# Dimensionality reduction reveals separate translation and rotation populations in the zebrafish hindbrain

**DOI:** 10.1101/2023.03.12.532285

**Authors:** Claudia E. Feierstein, Michelle H.M. de Goeij, Aaron D. Ostrovsky, Alexandre Laborde, Ruben Portugues, Michael B. Orger, Christian K. Machens

**Affiliations:** Champalimaud Neuroscience Programme, Champalimaud Foundation, Lisbon, 1400-038, Portugal; Faculty of Medicine, Utrecht University, Utrecht, 3584 CG, Netherlands; Institute of Neuroscience, Technical University, Munich, 80802, Germany; Munich Cluster of Systems Neurology (SyNergy), Munich, 81377, Germany

**Author notes:** Pfizer BV, Capelle aan den Ijssel, 2909 LD, Netherlands. Senior author, equal contribution (Michael B. Orger, Christian K. Machens).

## Abstract

In many brain areas, neuronal activity is associated with a variety of behavioral and environmental variables. In particular, neuronal responses in the zebrafish hindbrain relate to oculomotor and swimming variables, as well as sensory information. However, the precise functional organization of the neurons has been difficult to unravel since neuronal responses are heterogeneous. Here we used dimensionality reduction methods on neuronal population data to reveal the role of the hindbrain in visually-driven oculomotor behavior and swimming. We imaged neuronal activity in zebrafish expressing GCaMP6s in the nucleus of almost all neurons, while monitoring the behavioral response to gratings that rotated with different speeds. We then used reduced-rank regression, a method that condenses the sensory and motor variables into a smaller number of ‘features’, to predict the fluorescence traces of all ROIs (regions of interest). Despite the potential complexity of the visuo-motor transformation, our analysis revealed that the bulk of population activity can be explained by only two features. Based on the contribution of these features to each ROI’s activity, ROIs formed three clusters. One cluster was related to vergent movements and swimming, whereas the other two clusters related to leftward and rightward rotation. Voxels corresponding to these clusters were segregated anatomically, with leftward and rightward rotation clusters located selectively to the left and right hemispheres, respectively. Just as described in many cortical areas, our analysis revealed that single neuron complexity co-exists with a simpler population-level description, thereby providing insights into the organization of visuo-motor transformations in the hindbrain.

## Introduction

Many organisms exhibit behaviors that compensate for perceived self-motion. Among these behaviors are the optokinetic response^1–4^ (OKR), a conjugate movement of the eyes in response to whole-field rotation that helps to stabilize the retinal image, and the optomotor response^5–7^ (OMR), a directional movement of the whole body that helps the animal to remain at a fixed position and is observed in fish and flies.

Zebrafish display both OKR^3, 8^ and OMR^6, 7^ from an early age, and are therefore ideally suited for the study of these behaviors. In response to rotational optic flow, zebrafish execute conjugate movement of the eyes or turning bouts. In response to translational optic flow, zebrafish turn to orient themselves in the direction of motion and swim forward to cancel the optic flow. To engage in either behavior, fish need to disambiguate rotational and translational optic flow, and convert them into appropriate motor commands that compensate for the observed motion.

Horizontal eye movements are controlled by motoneurons residing in the oculomotor and abducens nuclei, which directly innervate the eye muscles, and are therefore the ultimate recipients of signals to control the eyes. Two main hypotheses have been proposed about the underlying motor circuitry for eye control. Hering’s ‘law of equal innervation’ argues for a dedicated circuit controlling conjugate eye movements, and also includes a circuit for vergent eye movements^9, 10^. Helmholtz’s theory instead proposes separate circuits controlling left and right eye movements, in which case conjugate (or vergent) eye movements arise through the circuits’ coordinated activation^11, 12^. Evidence for the two hypotheses has remained inconclusive: in the monkey brain, some studies have identified separate areas for conjugate and vergent eye movements^13, 14^, while others have pointed to monocular encoding of eye movements^15–17^. Similarly, studies in teleost fish find both monocular and binocular encoding of eye position and velocity in all areas of the oculomotor system that have so far been surveyed^18–20^.

Tail movements are controlled by spinal cord circuits, which receive inputs from different reticulospinal neuron groups that mediate forward swimming and turning bouts^21^. The anterior hindbrain, a region that is likely upstream of the reticulospinal system, has also been implicated in the selection of turning direction in different contexts, although it may not directly drive motor output^22–26^. Interestingly, this brain region is also active during spontaneous eye saccades, suggesting that this area could be coordinating eye and tail movement in this behavioral context^25^. Whether other hindbrain areas also mediate both tail and eye movements remains unknown.

While the studies described above have provided important insights into the functioning of these systems, they have largely focused on the analysis of single neurons, showing that neurons in the hindbrain are diversely tuned to monocular or binocular eye position and eye velocity, or correlated with directional swimming. Zebrafish are an ideal model organism to move beyond single-neuron analysis because they allow imaging of large parts of the brain, or even complete brains, at cellular resolution^25, 27–30^. Several studies have taken advantage of this, mapping behavioral variables onto large sets of individual neurons and performing functional clustering to identify brain areas or candidate circuits active in a range of behaviors ^22, 23, 25, 30–33^.

Still, these approaches do not solve the deeper problem of the puzzling tuning diversity of individual neurons. Heterogeneity of tuning, and specifically the mixing of both sensory and motor parameters, is a well-known property of many higher-order cortical circuits^34–36^, and can often be given a simpler and more intuitive interpretation when studied with methods that reduce the dimensionality of population activity^37–41^. Such methods replace the multitude of single-neuron tuning curves by a few population modes that summarize the activity of a whole population and capture its ‘essence’.

Here we set out to study the hindbrain, an area that has been strongly associated with both eye and tail movements^18, 22–25, 30, 32, 42^. The majority of studies of eye-related responses have focused on ventral areas, including the abducens nucleus and the oculomotor integrator. In a previous whole-brain study^30^, however, we identified a population of eye-sensitive neurons located in the dorsal hindbrain, and thus here we further explore this region. To capture a large range of behaviors, we used a set of visual stimuli aimed at dissociating movements of the left and right eyes, thereby eliciting a diversity of behavioral responses that include rotation and vergence eye movements, with accompanying swimming. Using two-photon Ca-imaging, we then recorded activity of tens of thousands of neurons while the fish were behaving. Through a combination of regression and dimensionality reduction, we find that the population activity in the hindbrain related to eye or tail movement is largely restricted to a two-dimensional subspace corresponding to rotation and vergence variables. Based on these dominant activity patterns, we show that neurons in the hindbrain can be separated into three anatomical segregated populations, one active during vergent motion and swimming, and two that are active during left or right rotational motion and swimming. Our work provides new insights into the organization of hindbrain circuitry and suggests a solution to the longstanding debate about circuits dedicated to vergent and rotational movements.

## Results

### Uncorrelated motion stimuli cause swimming bouts and decoupled eye movements

Larvae were presented with motion stimuli to induce horizontal eye rotations, a reflexive behavior in response to whole-field rotation. Stimuli consisted of a rotating striped pattern projected onto a screen below the head-restrained fish. The rotation of the grating was modulated with steps of constant velocity (see Methods; Figure 1A), in order to decouple eye position and velocity. In addition, to decouple movement of the left and right eyes, we divided the screen into two hemifields, and the velocity of the projected gratings was modulated independently in the left and right hemifields (Figure 1A and Figure S1A; Movie S1). In this way, stimuli delivered to the two eyes were, on average, uncorrelated. Five different grating velocities were presented to each eye (-30, -15, 0, 15 and 30 °/s), resulting in 25 different stimulus combinations; each of these combinations was presented for five seconds. The 25 stimulus combinations formed a stimulus set that was repeated in each stimulus plane, in a pseudo-random order.

**Figure 1.**
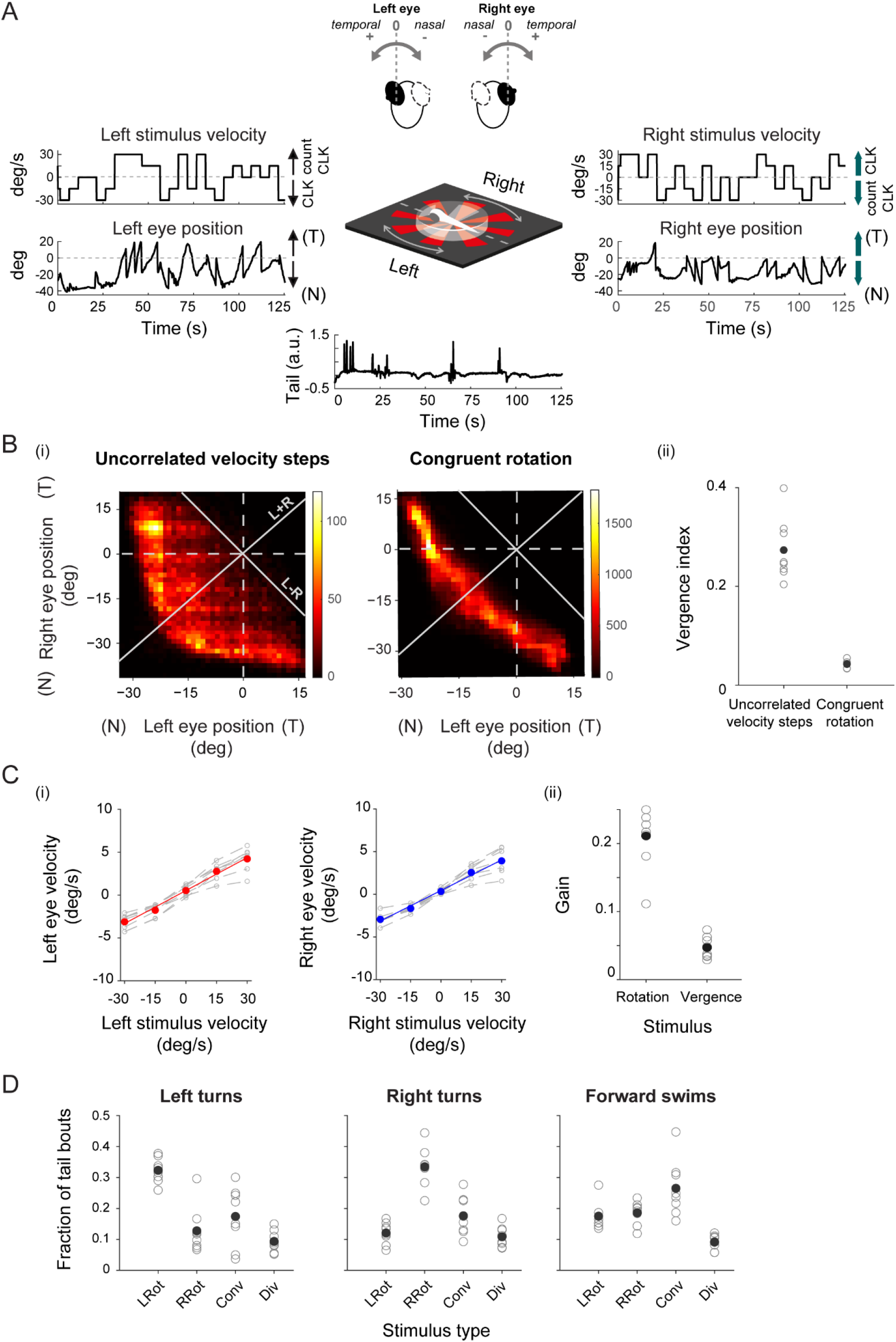
Split motion stimuli are used to decouple eye movements. **(A)** Schematic of visual stimulation (center). Eye rotations are elicited by a rotating radial pattern projected onto a screen placed below the fish. The projected stimulus is split in two, and the central, binocular area is devoid of stimulus (see Methods and Movie S1). The velocity of the presented pattern is modulated with steps of constant velocity. Note that each hemifield is modulated independently. Both eye and tail positions are tracked during stimulus viewing. **(B)** (i) Left and right eye positions are decorrelated when using a split stimulus (left, uncorrelated velocity steps). In comparison, left and right eye positions are highly correlated when using a regular optokinetic stimulus, i.e., a rotating grating with the same velocity for both eyes (right, congruent rotation). Eye positions can also be expressed as the sum or difference of the two eyes (white axes), which correspond to vergence and rotation of the eyes, respectively. (ii) Vergence index (variance of eye vergence / variance of eye rotation; see Methods) for fish presented with split, incongruent motion stimuli (8 fish) and congruent rotation stimuli (5 fish). Split stimuli elicit more vergent eye movements, as compared to congruent eye stimulation. Black filled circles, mean across fish. Open circles, individual fish. **(C)** Eye velocity is modulated by stimulus velocity (see also Figure S1). (i) Left eye and right eye velocities for each stimulus velocity presented on the corresponding half of the screen. Red and blue points correspond to the mean across fish (N=8), while gray traces correspond to individual fish. The slope of the fitted lines reflects the eye velocity gain, i.e., the modulation of eye velocity by stimulus velocity. (ii) Eye velocity gain, pooled for both eyes, for stimuli grouped into rotating and vergent. Filled circles, mean across fish (N=8). Open circles, individual fish (see Methods and Figure S1D). **(D)** Swimming bout type is modulated by stimulus category (see also Figure S1A). Left turns occur more frequently when the stimulus moves leftwards (counterclockwise); conversely, right turns are observed more frequently when the stimulus moves rightwards (clockwise). Forward swims are primarily observed when both stimuli move in the nasal direction, mimicking forward motion (Converging; see Methods and Figure S1E). Filled circles, mean across fish (N=8). Open circles, individual fish. Conv: converging. Div: diverging. LRot: leftwards rotation (counterclockwise). RRot: rightwards rotation (clockwise).

While the fish were presented with the stimuli, we simultaneously tracked eye and tail behavior at 100 and 700 Hz, respectively, using a high-speed, infrared-sensitive camera. Eye and tail positions were extracted, and eye velocities were estimated from changes of the eye positions (see Methods; Figure 1A).

As intended, the use of uncorrelated stimuli resulted in a partial decoupling of left and right eye movements (Figure 1Bi; Figure S1C). Eye positions thereby spanned a richer position-space, as compared to the classical optokinetic response (OKR), in which both eyes move congruently in response to a whole-field rotating stimulus (Figure 1Bi, right panel).

Stimuli, eye positions, and eye velocities can be described along ‘vergence’ and ‘rotation’ axes, which result from adding or subtracting the corresponding left and right variables (white axes in Figure 1Bi; see Methods). Along the vergence axis, stimulus gratings on both halves are then moving either forward (converging) or backwards (diverging), thereby mimicking translational motion. Similarly, eye positions and velocities are converging or diverging. Along the rotation axis, stimulus gratings on both halves are moving either towards the left (counterclockwise), or towards the right (clockwise), thereby mimicking rotational motion (Figure S1A). Similarly, eyes move conjugately either in clockwise or counterclockwise directions.

Whereas congruently rotating stimuli drove eyes to rotate (note that eye positions lie on a line parallel to the rotation axis; Figure 1Bi, right panel), using uncorrelated velocity steps drove the eyes to converging (both eyes directed nasally) as well as diverging (both eyes directed temporally) positions (Figure 1Bi-1Bii). To quantify the difference in prevalence of vergence positions, we defined a ‘vergence index’ as the ratio of the variance of vergent eye positions over the variance of rotation eye positions (Figure 1Bii). This index was higher for all fish presented with incongruently rotating gratings, as compared with fish presented with congruently rotating gratings. Convergent eye positions, usually observed during prey capture^43^, were readily driven with the stimuli used here, namely during the presentation of back-to-front stimuli to both eyes. Divergent eye positions were, on the other hand, underrepresented; this is consistent with the observation that fish species with yoked eye movements show reduced or absent responses to front-to-back whole-field motion^44, 45^. The velocity of either eye increased linearly with the velocity of the stimulus in the corresponding hemifield (Figure 1Ci; Figure S1B-C). However, the gain with which eye movements were modulated was not equal across stimuli: Fish tracked rotating stimuli with higher gain than translational (vergent) stimuli (Figure 1Cii; Figure S1D).

Eye movements were accompanied by tail flicks, which could be classified as left and right turns, and forward swims^32^ (see Methods). Forward swims occurred mostly when fish were presented with vergent stimuli moving nasally (converging), which resemble forward translational flow (Figure 1D; Figure S1A,E). Turns occurred less frequently (not shown), but their directionality was determined by the stimulus: overall, left rotating stimuli drove turns to the left, while right rotating stimuli drove turns to the right (Figure 1D; Figure S1E).

### Hindbrain activity is highly correlated with eye and stimulus variables

To understand the role of the hindbrain in stimulus-driven oculomotor behavior, we recorded hindbrain activity in six-to seven-day-old larvae expressing GCaMP6s in the nuclei of most neurons^46^ (*elavl3*:H2B-GCaMP6s), using a two-photon microscope, with an imaging rate of 2 Hz (see Methods; Figure 2A-D). We simultaneously presented the uncorrelated motion stimuli and monitored the larval behavioral response. We targeted volumes (width x length x depth) of 369μm x 246μm x 80μm up to 369μm x 246μm x 110μm of the hindbrain, which spanned the dorsal aspect of the intermediate and inferior medulla oblongata, situated in rhombomeres 2 to 7. For each larva (N=8 larvae), we imaged the volume in the dorsal hindbrain as a stack of 80-110 planes in 1μm steps. For each plane, the 25 stimulus combinations were presented in a randomized order, before advancing to the next plane. Fish showed sustained behavioral responses throughout the imaging session (Figure S1B).

**Figure 2.**
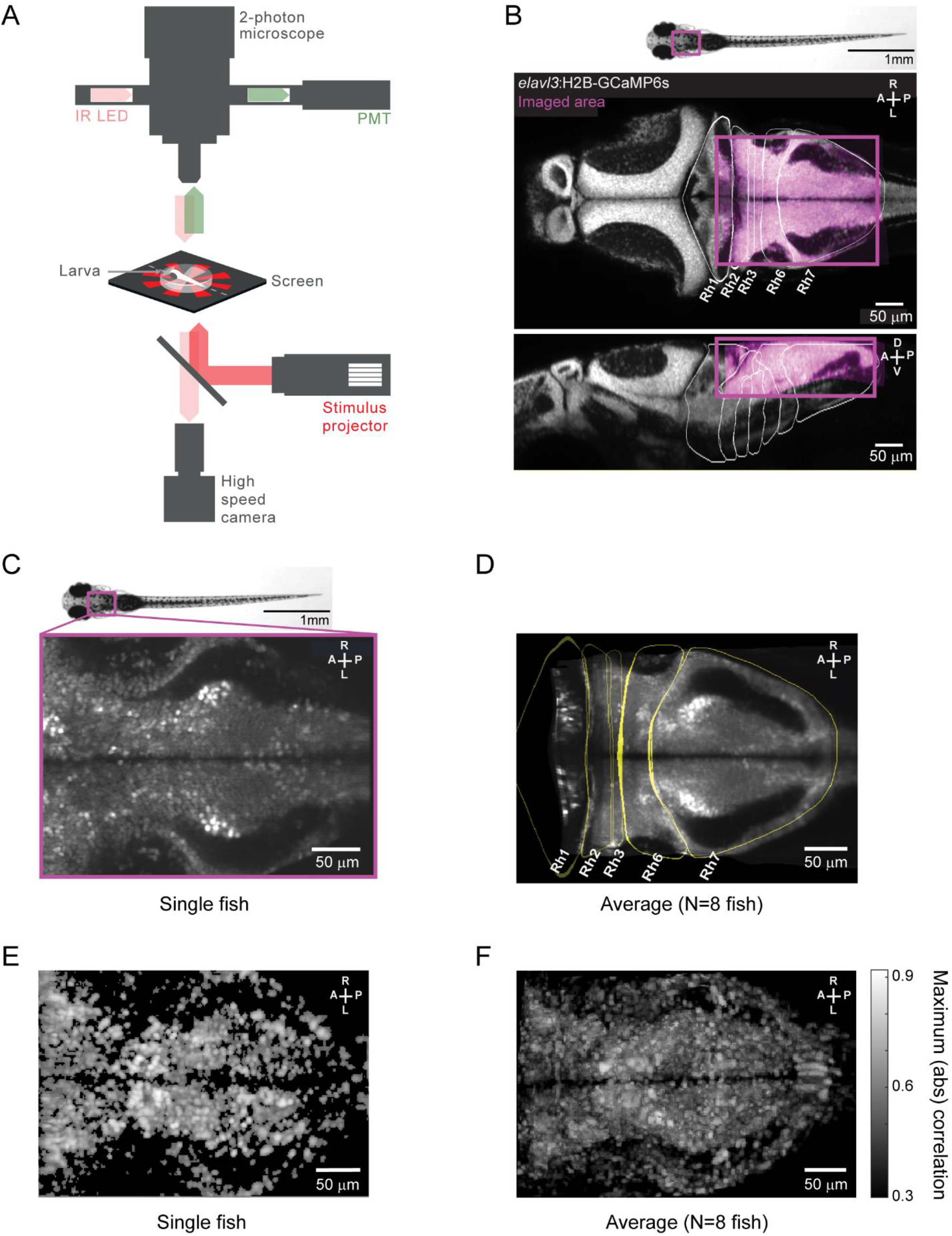
Activity related to behavior is widespread in the zebrafish hindbrain. **(A)** We imaged the hindbrain using a two-photon microscope during split-screen motion stimulation. Schematic of experimental setup (see Methods), showing simultaneous brain imaging, behavior monitoring and visual stimulus presentation. **(B)** Representative plane highlighting the imaged region (magenta) overlaid on the corresponding plane of the Z-brain atlas^47^. Top, dorsal view. Bottom, sagittal view. **(C)** Example plane of the imaged hindbrain area in a representative fish. **(D)** Example plane of the average image stack across eight fish. Hindbrains were registered to an internal template (see Methods) and then to the Z-brain atlas, before averaging. **(E)** Maximum absolute correlation value with any of the expanded set of behavioral regressors (see Methods) for an example fish (maximum projection of the imaged stack). The resulting map was registered to the Z-brain atlas. **(F)** Average maximum absolute correlation maps across eight fish, after registration to an internal template (see Methods; maximum projection of average stack). Correlation maps are thresholded at 0.3 absolute correlation value.

Activity, measured as fluorescence changes, was extracted from the imaging data after preprocessing (see Methods). Briefly, data was first motion-corrected, and then segmented by dividing each hindbrain stack in cuboid-shaped regions-of-interest (ROIs). The resulting ROIs were overlapping and tiled the whole imaged area. We then sought to characterize the relationship between the neural activity and various experimental and behavioral variables, including eye position and velocity, stimulus velocity, and tail behavior. The complete set of these 12 ‘original’ variables is shown in Figure S2.

Neuronal activity could, in principle, be explained not only by the *present* state of the behavior/stimulus variables, but could also depend on their *past* or *future* state (the latter, for instance, for activity that would predict a motor command). Indeed, the dynamics of calcium indicator alone introduces a dependence of current neuronal activity onto future measurements. To account for any dependencies on past or future states, we built a set of additional ‘shifted’ variables: each original variable was shifted by up to 20 seconds into the past and 10 seconds into the future (see Methods).

We first correlated activity to our regressors as well as various thresholded variants and combinations of them (see Methods and Table 1). This approach has proven useful to reveal areas active during a given behavior^26, 30, 42^. Regressor-activity correlations were widespread in the hindbrain: A large population of ROIs (voxels) was highly correlated with at least one of the stimulus and behavioral variables included in the analysis (Figures 2E-F).

Correlation analysis provides a first overview of how this brain area may be related to sensory and motor information. However, we find that many individual ROIs correlate with *both* sensory and motor variables (not shown; see also ref. 30). Since regressors are correlated with each other to some degree, this mixing of sensory and motor variables in neural activities may simply reflect such latent correlations, rather than a true correlation of the neural activities with the respective sensory and motor variables. To investigate how ROI activity could be explained by our set of regressors, we turned to multiple linear regression (MLR).

### Hindbrain activity can be explained by a low-dimensional set of features

Classical MLR finds for each ROI in the regression model a set of weights that explains the activity of that ROI as a function of the regressors (Figure 3A, S3A). As described above, our shifted regressors covered time windows in the past and future. We reasoned that this approach would capture the delay introduced by the calcium indicator, without making any assumptions about the dynamics of the calcium kernel. For a given ROI, we can then find a linear combination of the shifted regressors, that is, a pattern of weights, that explains the activity of the ROI as a function of the original variables and their time shifts (Figure 3A). The shifted weights of each original variable can be represented as a kernel (see inset in Figure 3A), where negative shifts reflect how past states of the variable influence activity, whereas positive shifts reflect how future states of the variable influence activity (the zero time-shift corresponds to the state of the variable at the time the activity was measured).

**Figure 3.**
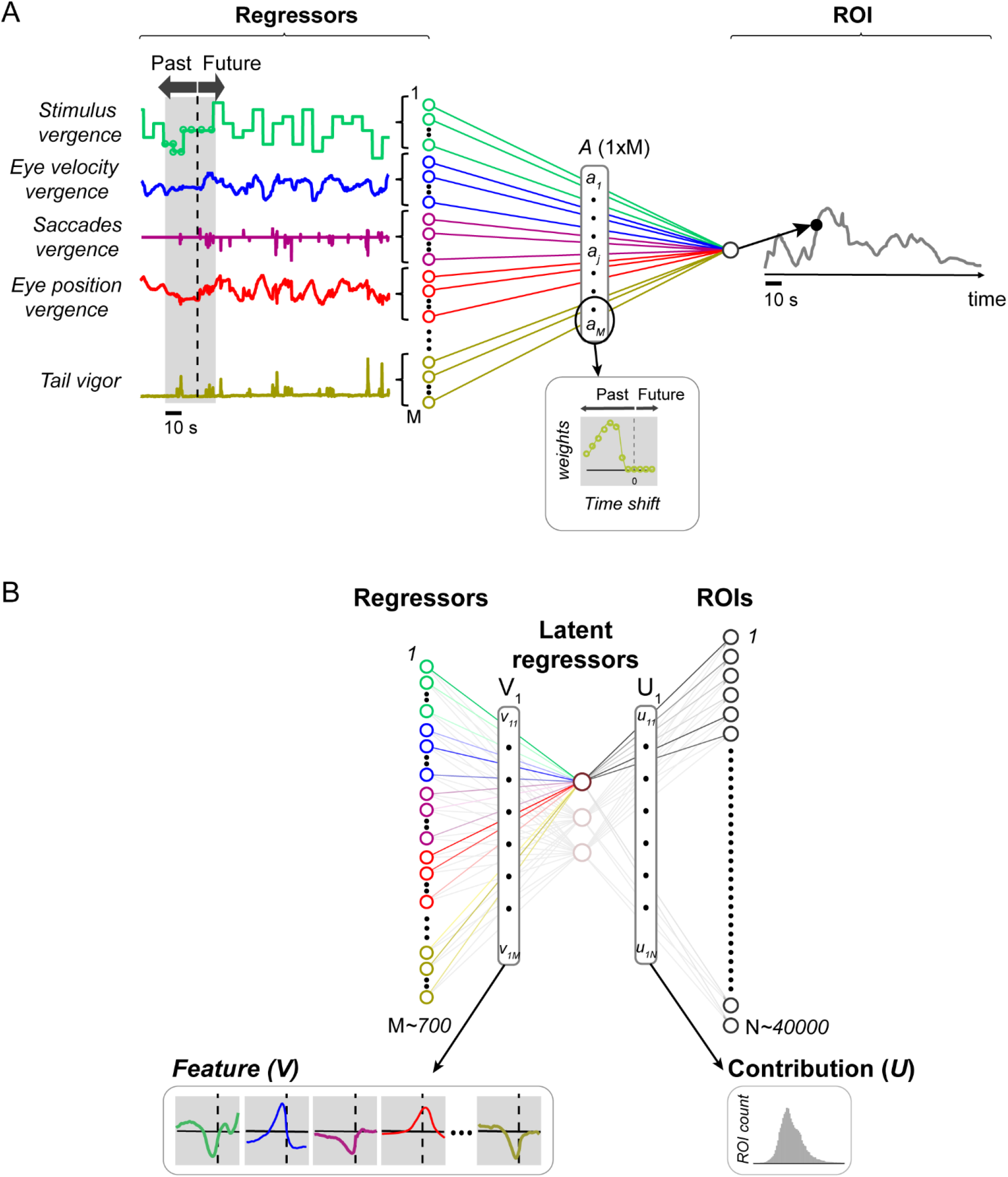
Regression models. **(A)** Multiple linear regression (MLR) model for a single ROI, using regressors in time windows. To account for activity-behavior relations that are not limited to the present time point, we expanded each regressor or behavioral variable (only a subset shown here) 20 seconds into the past and 10 seconds into the future. In this way, for each original variable (see Figure S2), the regression model finds a set of weights that corresponds to the contribution of the variable at different time points, to ROI activity at the present time point. The vector of regression weights, *A*, includes these time-shifts. Inset shows the weights for different time points of one of the variables. Dotted line corresponds to no time-shift. A feature (highlighted) corresponds to a pattern of weights for the regressors (V_1_, in this example). The weighted combination of the original regressors then gives rise to a latent regressor. In turn, this latent regressor is associated with a pattern of ROI activity, determined by its contribution weights (U_1_). Left inset shows the weights of each time point for the different regressors, as in (A) (each regressor is represented in one color). Right inset shows distribution of the U values (contribution) for this example feature. This distribution illustrates how strongly the latent regressor associated with this feature is expressed across the population.

However, this approach has two downsides. First, by analyzing each ROI individually, we are not truly leveraging the size of the data to discover features or activation patterns spread across the whole population. Second, it is not immediately clear how to interpret all of the weights found by MLR in a succinct manner. We therefore turned to reduced-rank regression^48^ (RRR), a variant of multiple linear regression which enforces a rank constraint on the matrix of weights and has proven useful in the analysis of neural populations^39, 49, 50^. Whereas classical MLR finds for each ROI a separate mapping from the regressors to the ROI activity (Figure 3A, Figure S3A), RRR first finds a low-dimensional subspace of *latent* regressors, which can be thought of as a small set of latent variables, and then finds, for each ROI, a separate mapping from the latent regressors to the ROI activity (see Methods; Figure 3B, Figure S3B). Formally, the latent regressors (or latent variables) are linear combinations of the original regressors, and we will refer to the respective combination of weights as *features*. In turn, the activity of each ROI is explained by a specific, linear combination of these latent regressors, and we will refer to these weights as the *contributions* of the latent regressors to the ROIs (Figure S3B; Figure 3B). Thus, a feature corresponds to a pattern of regressors that, whenever present, gives rise to a particular pattern of ROI activities (Figure 3B). Since our regressors are time-shifted, each feature is composed of a set of ‘kernels’ (inset in Figure 3B), and each kernel contains the weights of one original regressor at different time points in the window analyzed.

We built separate RRR (as well as MLR) models for eight different fish, including 732 regressors (12 original regressors plus their shifts) and 26000 to 57000 ROIs, depending on the fish (median 45588 ROIs/fish; see Methods). The RRR models were regularized and cross-validated (see Methods; Figure S5A). Figure 4 shows the results for an example fish. For this fish, we found that the best model had rank six, i.e., six latent regressors were sufficient to explain the activity of a population of 43622 ROIs. The performance of the selected model, measured as explained variance, was estimated for each individual ROI (see Methods) and is shown in Figure 4A. The mean explained variance in the test set was 0.46, with some ROIs having explained variance as high as 0.76 (Figure 4A, S6D; note that the mean predictive power depends on the criterion used to include ROIs in the model; see Methods).

**Figure 4.**
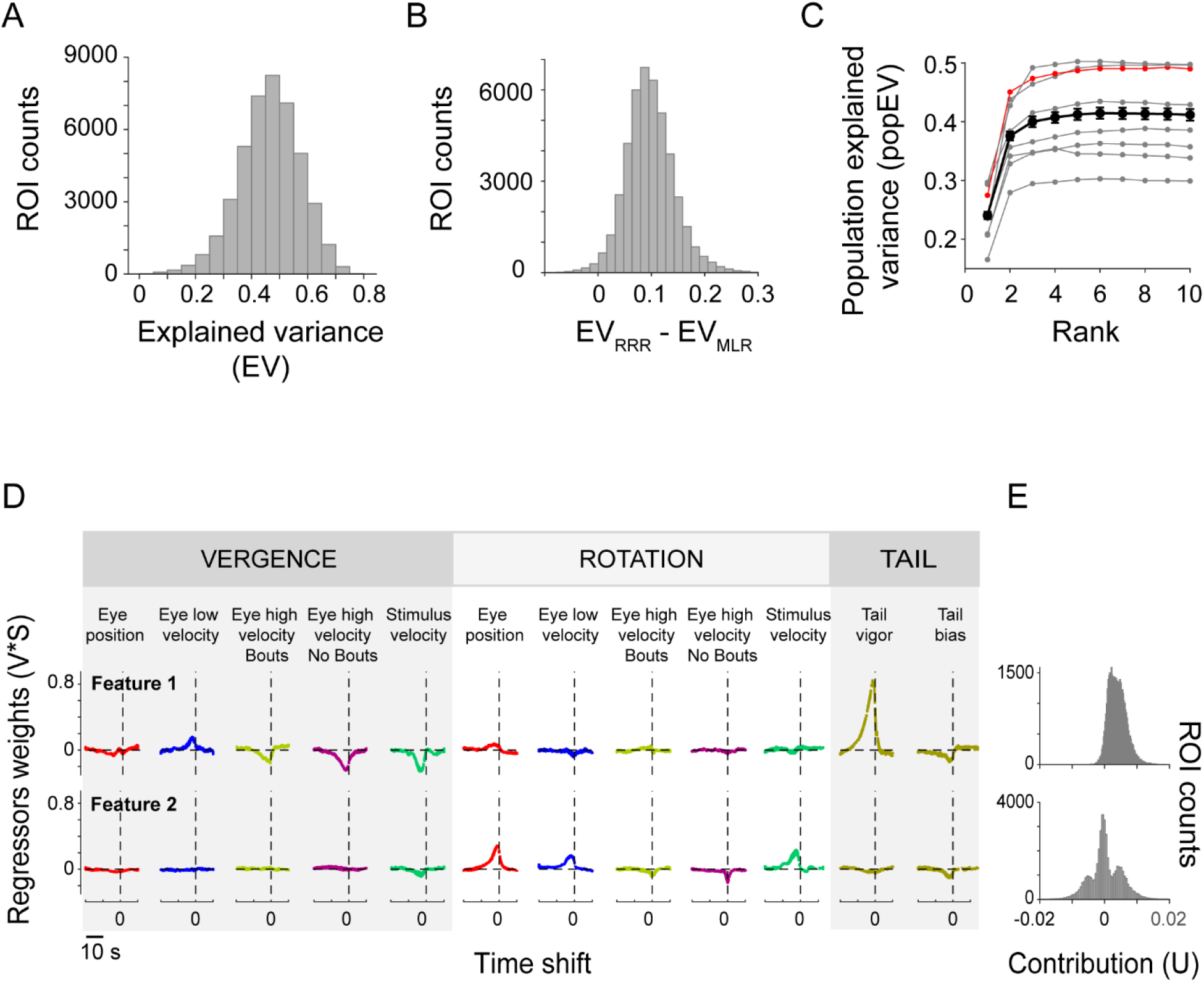
Activity is explained by a small number of features that separate vergence and rotation variables. **(A)** Explained variance (EV; cross-validated) in the test data for the ROIs of an example fish using RRR. **(B)** The explained variance of the best RRR model (EV_RRR_) was generally higher than the explained variance of the regular multiple linear regression model (EV_MLR_). Same example fish as in (A). Population explained variance (popEV; see Methods) as a function of the number of features (rank) included in the model for each individual fish (gray traces; example fish in A,B is shown in red) and across fish (average 8 fish, black trace). In most fish, the first two or three features contribute most of the explainable variance. **(D)** The first two features for the example fish. Feature traces are scaled by their overall importance (V*S; see Methods). For each feature, variables are color-coded and sorted into vergence, rotation, and tail variables. Vertical dotted line highlights a time shift of zero. Traces of five cross-validation runs are shown overlaid. Note that feature 1 groups activity related to vergence and swimming, whereas feature 2 groups activity related to rotation. Features 3-6 (see Figure S6) of this fish are also statistically significant, but they only have a minor contribution to the explained variance. **(E)** Contribution of the latent regressors associated with features 1 (vergence) and 2 (rotation) to ROI activity for the same example fish (N= 43622 ROIs).

We next asked whether the dimensionality reduction step inherent to RRR gains explanatory power for some ROIs at the cost of losing explanatory power for others. Such a tradeoff could occur when the features, which capture only a subspace of the full regressor space, are ill-matched for some (outlier) ROIs. We therefore compared the performance of the RRR and the full MLR models for individual ROIs, using the same regularization parameter. For almost all ROIs, predictive power was higher for the RRR model than the MLR model (Figure 4B) and we found no evidence for clear outliers. Thus, the latent regressors found by the RRR model capture as much variance as possible given the visual stimuli and behavioral variables included.

For the example fish, the first two latent regressors contributed to almost all (92%) of the population explained variance. Indeed, even though across fish the best models had between three and six latent regressors, on average, only little explanatory power was gained by the model when including more than two latent regressors (Figure 4C; red curve corresponds to the example fish). Using RRR, we therefore greatly shrank the complexity of the problem, and revealed an effectively two-dimensional subspace of regressors that captures the bulk of hindbrain population activity. This strong drop in dimensionality was caused by correlations among the ROIs rather than correlations among the regressors, as the regressor space had many more dimensions (Figures S4C,D). For the rest of the analysis, we therefore focused on the first two latent regressors.

### Hindbrain activity is largely explained by rotation and vergence features

As described above, each latent regressor is generated through a pattern of regressor weights, or a feature (Figure 3B, Figure S3B). Figure 4D shows the first two features for the example fish (all six features of the model can be found in Figure S6). The first feature is almost exclusively composed of vergence regressors (Figure 4D, top row). These regressors include the stimulus vergence (translational motion), and the vergent eye velocities. In addition, the first feature also consists of tail vigor, which essentially measures forward swimming. This feature therefore combines sensory and motor information. The second feature, on the other hand, is exclusively made of rotation regressors again including sensory (stimulus rotation) and motor information (rotation of the eyes, both in terms of eye position and eye velocity). Thus, the two features which contribute the most to explaining population activity in our model, seem to separate vergence and rotation information. In what follows, we will refer to them as *vergence* and *rotation* features, respectively. We note that saccade-related activity was only observed in the vergence feature, and that tail bias, which reflects the laterality of the tail flick, was present in both features, with a small contribution.

When we built models for eight different fish, we found similar vergence and rotation features in all of them (not shown). In six out of eight fish, the vergence and rotation features corresponded to the first two features; in the two remaining cases, vergence and rotation features could be identified amongst the first three features of the model.

In other words, our analysis shows that there are patterns in the population activity that are related to vergence in the stimulus and behavioral variables, namely stimulus vergence, eye velocity vergence and swimming, and there are separate patterns of activity that are related to rotation in the stimulus and behavioral variables, namely stimulus rotation and rotation of the eyes.

We also studied the relative importance of the 12 original regressors in explaining neuronal activity. To do so, we turned to variable selection. We first performed forward stepwise variable selection^51^, where models are grown one variable at a time (in our case, an initial variable with its shifts; Figure S7A right panel and S7B). The most important regressor was tail vigor, which accounted on average for 50% of the variance explained by the full model. The second place went to a rotational variable, either eye position rotation (4/8 fish), stimulus velocity rotation (2/8 fish), or eye velocity rotation (1/8 fish), and yielded on average, 84% of the explainable variance. These latter regressors were to a certain extent exchangeable. The third-largest effect was, most often, given by the vergence of the stimulus velocity, bringing the count up to 91% of the explainable variance. We next tried backwards stepwise selection ^51^, where variables are removed one at a time. As with the forward approach, tail vigor was the most important variable, as removing it produced the biggest drop in model performance (Figure S7A, left panel). The second place went for eye position rotation, although the decrease in model performance was much smaller when eliminating this variable. Removing the rest of the variables resulted in no loss in the amount of variance explained by the model, suggesting that these variables are likely redundant.

### ROIs cluster into three groups in feature space

We next asked how the latent regressors contribute to the activity of each ROI by plotting the distributions of their contribution values U (see Methods; Figure 3B). The latent regressor associated with the vergence feature showed an asymmetric contribution to ROI activity: for the vast majority of ROIs, this latent regressor had a positive contribution to activity (Figure 4E, top row). The latent regressor associated with the rotation feature, on the other hand, had a symmetric contribution to ROI activity: for some ROIs, activity was positively related to this latent regressor, while for others it was negatively related (Figure 4E, middle row), suggesting that different ROIs are active during leftwards or rightwards rotations, respectively.

It is possible that each ROI expresses a random combination of the latent regressors associated with vergence and rotation; alternatively, the latent vergence and rotation regressors could be expressed in separate sets of ROIs, e.g., corresponding to separate anatomical populations. When looking at the joint contribution of the latent vergence and rotation regressors to ROI activity for our example fish (Figure 5A), it becomes apparent that ROIs form three main groups. We therefore assigned ROIs to clusters using either manual or automatic clustering methods. In Figure 5B, we drew a polygon around the clusters, and then plotted in Figure 5C the ROIs color-coded according to their cluster assignment. Using unsupervised clustering in multiple dimensions (ClusterDv^52^), the same three clusters were found (Figure 5D). The asymmetry we observed in the contribution of the latent vergence regressor is also evident here: all clusters have a positive contribution of the latent vergence regressor.

**Figure 5.**
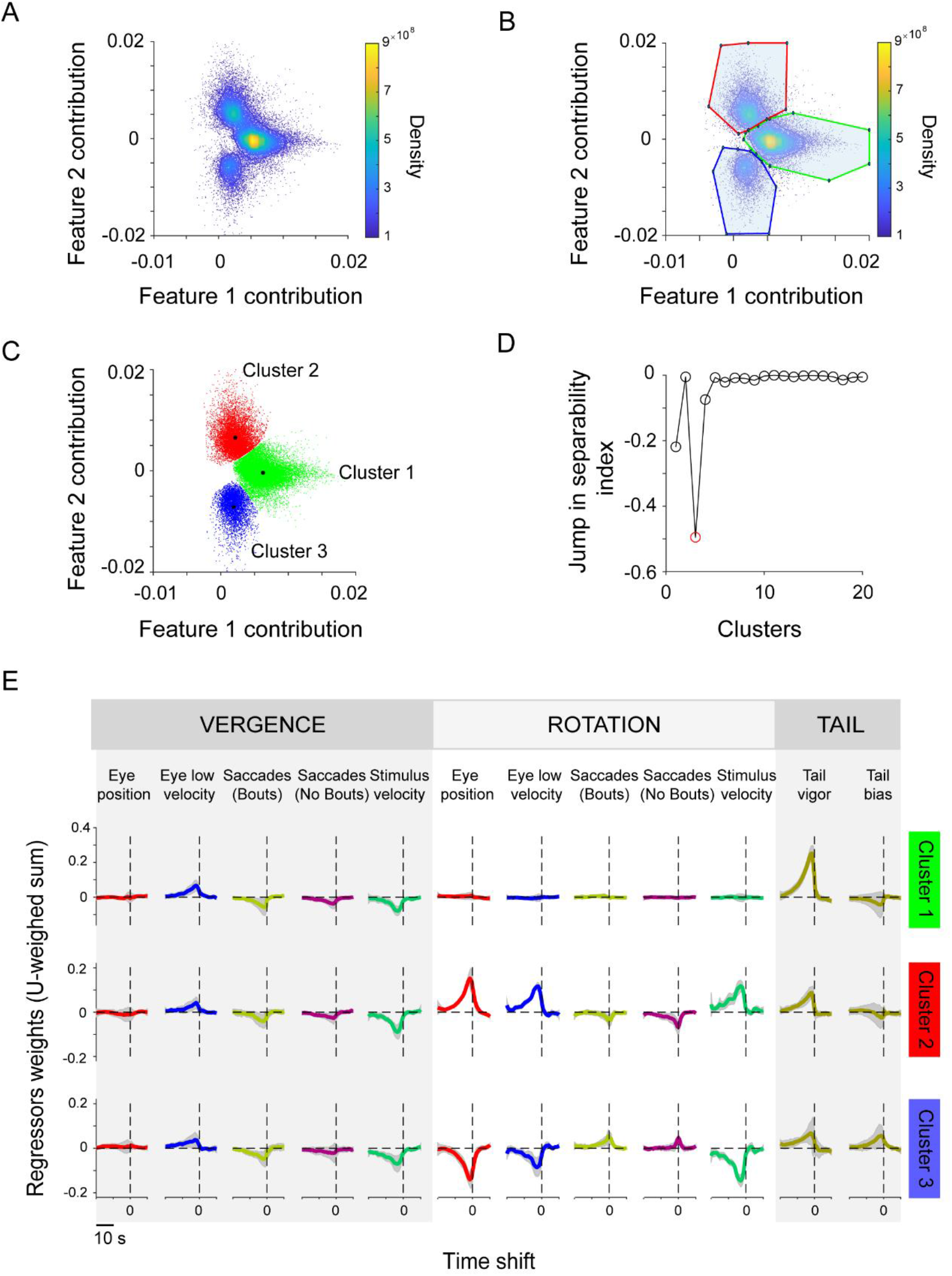
ROIs form three clusters that relate to vergence and left/right rotational motion. **(A)** ROIs can be grouped into three main clusters based on the contribution of vergence and rotation features to their activity. Contribution (U) of latent regressors associated with features 1 (vergence) and 2 (rotation) to ROI activity. Each dot corresponds to an ROI in an example fish (N=32228 ROIs with EV>0.4). Three clusters become apparent. **(B)** Manual assignment of ROIs to three clusters in feature contribution space (U1-U2; see Methods) based on the density plot. **(C)** ROIs for the example fish are color-coded according to their cluster assignment (ROIs are subsampled here for better visualization). Black asterisks mark cluster centroids. **(D)** Using ClusterDv^52^ (see Methods), we also find three clusters. **(E)** Clusters relate to vergence and left/right rotational motion. The weighted contribution of features 1 to 3 is shown each cluster’s centroid. Each row corresponds to a cluster, and variables are grouped into vergence, rotation and tail variables (compare to Figure 4D). Cluster 1 (green) is composed exclusively of vergence variables (note that vergent stimuli can be interpreted as translational motion) and swimming. Clusters 2 (red) and 3 (blue) relate to leftward and rightward rotation, respectively. Traces are mean±STD across 7 fish (one fish was excluded here because the tail tracking was too noisy).

We performed clustering in all other fish in our data set, and found that these three main clusters appear in all examined fish (Figure S8). Note that in some fish, these clusters can likely be further subdivided: in at least two of the fish (fish 6 and 8) one can see a further subdivision of one of the clusters; this was not further investigated, as the clustering algorithm was unable to separate them. Thus, ROIs in the hindbrain form three principal groups according to how the latent vergence and rotation regressors contribute to their activity.

### The three clusters represent vergence, leftward rotation, and rightward rotation

Within each cluster, individual ROIs exhibited various contributions of the two latent regressors. This observation held particularly for cluster 2 (red) and cluster 3 (blue), in which ROIs had a strong contribution of the latent rotation regressor, but also a smaller, positive contribution of the latent vergence regressor.

To understand what each of these three main clusters represent, we examined a representative point of each cluster, its centroid. For each centroid, we can then visualize the regressor weights that result from combining the relevant features (Figure 5D). Cluster 1 (green) contains mostly contributions of vergent variables, including stimulus velocity and eye velocity, as well as swimming. ROIs in this cluster will be active whenever the fish is presented with vergent stimuli and engaged in a vergent movement of the eyes and swimming. We will refer therefore to this cluster as the *vergence cluster*. Because stimulus and (low) eye velocity appear with opposite signs, consistent with a representation of retinal slip, we hypothesize that ROIs in this cluster could represent information related to this variable, or some other function of translational flow. Interestingly, convergent (forward translation flow) stimuli elicit swimming (see Figure 1D), which is also represented in this cluster (see tail vigor).

Clusters 2 (red) and 3 (blue), on the other hand, are dominated by rotational variables, suggesting that ROIs in these clusters could represent an estimate of rotational flow, either counterclockwise (red cluster, positive contribution of rotation variables) or clockwise (blue cluster, negative contribution of rotation variables). We will therefore refer to these clusters as the *leftwards (counterclockwise) rotation cluster* and *rightwards (clockwise) rotation clusters*, respectively. We note that the contribution of tail bias (whose strength was somewhat variable across fish) was consistent with the behavioral observations: leftwards rotating stimuli elicited preferably a left turn (Figure 1E). Accordingly, ROIs active during leftward rotations (leftwards rotation cluster, red) showed activity related to left turns (note that a negative bias corresponds to a left turn; see Methods). The reverse is true for rightward turns.

Although the left and right rotation clusters are dominated by rotation variables, they also include vergence variables. Importantly, the spread of ROIs within each rotation cluster shows that different ROIs have different degrees of contribution of the vergence (feature 1) variables (Figure 5C). For eye and stimulus velocity, this resulted in a continuum that ranged from ROIs representing bilateral eye and stimulus velocity rotation, to ROIs representing ipsilateral eye, yet contralateral stimulus velocity (albeit in ipsiversive direction; Figure S9). In other words, ROIs combined both eyes and represented rotation, but mixed ipsilateral and contralateral sensory and motor variables in counterintuitive ways. The representation of eye position was simpler and did not change much across the cluster, with both clusters representing conjugate eye position. (We note that our imaging area did not cover the oculomotor integrator or the abducens).

Monocular and binocular encoding of saccades^20^ and eye velocity^18^ has recently been described in zebrafish larvae. Although in the latter study a spatial gradient of binocular/monocular activity was described, we did not find such a clear spatial organization (data not shown). This discrepancy may arise from differences in the imaged brain areas.

### ROIs in different clusters correspond to separate anatomical populations

To visualize the anatomical distribution of ROIs belonging to these clusters, we built maps where we color-coded each ROI according to its cluster membership (Figure 6; Figure S10). ROIs in the vergence cluster were colored green, and ROIs in the leftwards and rightwards rotation clusters were colored in red and blue, respectively. Clusters were spatially segregated, with the rightwards and leftwards rotation clusters symmetrically localized to the right and left hemispheres, respectively, while the vergence cluster was distributed in both hemispheres. The rotation clusters were distributed in more compact areas, whereas ROIs belonging to the vergence cluster were more widely distributed. Because ROIs in the borders of the cluster may have not been properly assigned (especially when there is greater overlap between points), we also examined the maps resulting from restricting the ROIs used to those within an ellipse around the centroid (in clustering space). The resulting maps (not shown) were qualitatively indistinguishable to the ones shown here, which also further supports the validity of the original clustering. Across fish, the anatomical organization of the clusters was similar (Figure 6; Figure S10; Movie S2).

**Figure 6.**
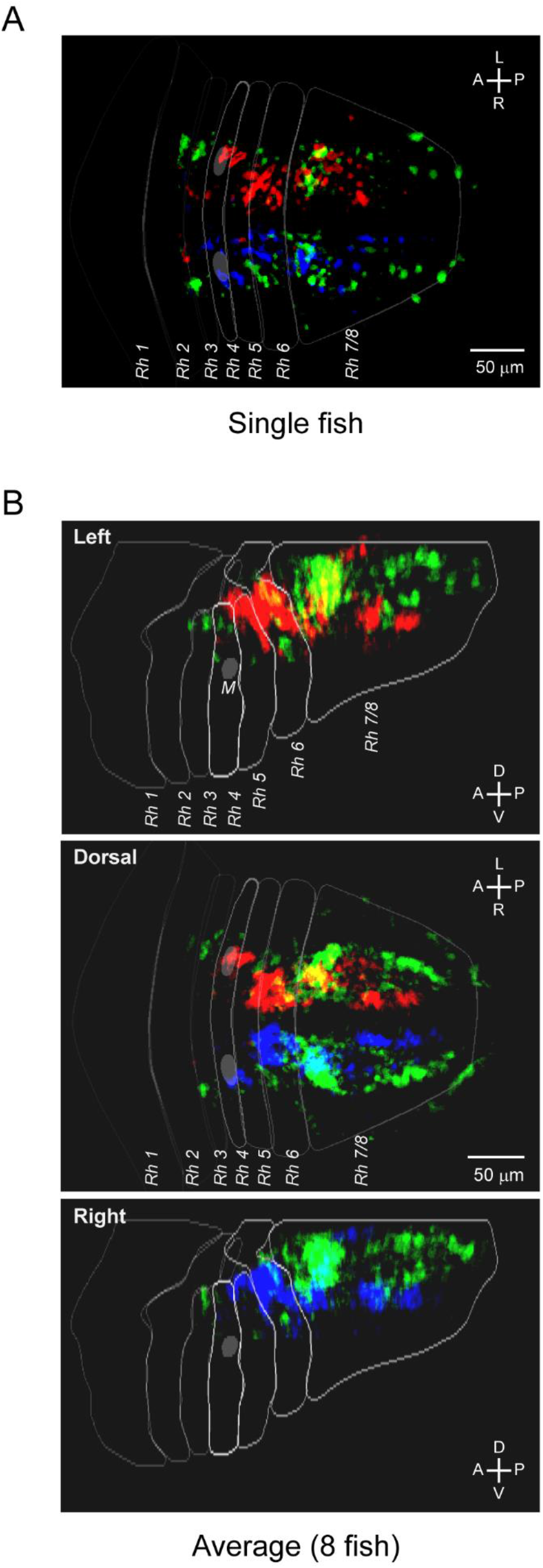
Anatomical distribution of ROIs belonging to the vergence and rotation clusters. **(A)** Spatial distribution of the ROIs assigned to the three clusters in the example fish (N=32228 ROIs with EV>0.4). **(B)** Average distribution of the ROIs assigned to the three clusters (N=8 fish). Individual maps (see Figure S10) were registered to the Z-brain template before averaging. Scale bar, 50 microns. Dark gray ellipse shows the location of the Mauthner cells (M). Rhombomere contours and Mauthner cell correspond to Z-brain masks^47^.

The three clusters spanned the whole imaged area, which corresponded to the dorsal aspect of the intermediate and inferior medulla oblongata. Previous studies^18, 42, 53^ looking at eye position and velocity encoding have focused on more ventral areas of the hindbrain, and thus, mostly non-overlapping with areas imaged in the present study. In a recent study^20^, the authors reported neurons active during saccadic eye movements located in rhombomeres 5 and 6; these neurons may partly overlap with our recorded region (see Discussion). The clusters defined here did not clearly overlap with any of the annotated hindbrain nuclei (MPI2, Z-brain), except for some overlap with the facial motor nucleus, and some overlap of ROIs in the green (vergence) cluster with the medial octavolateralis nucleus, and more prominently with the vagal sensory lobe and vagus motor nucleus (MPI2 atlas). In addition, we did find some overlap with some molecularly defined areas. The vergence cluster overlapped with neurons expressing hypocretin receptor (6.7FDhcrtR-Gal4 Stripe 1, Z-brain), neurons expressing Gad1b (Gad1b stripes 1 and 2, Z-brain), and more prominently, neurons expressing Ptf1a (Ptf1a Stripe, Z-brain). Neurons in the rotation clusters also overlapped with a population expressing the hypocretin receptor (6.7FDhcrtR-Gal4 Cluster 1, Z-brain). Neurons in these clusters also mapped to regions expressing Gad1b (Gad1b stripe 3, Z-brain), and more prominently to areas expressing the glycinergic transporter (Glyt2 cluster 1, and stripes 1-3, Z-brain), Otpb and Ptf1a. These anatomical overlaps provide some hints to the neurotransmitter identity of the neurons belonging to our functional clusters, and future experiments should further investigate the involvement of different neuronal types in this behavior.

Thus, our analysis shows that, during rotational and vergence eye movements, two different populations in the dorsal medulla are active. One population relates to translational motion, while the other to rotational motion. Similar to other findings^18^, neurons responding during rotational motion showed a continuum of responses, ranging from neurons representing binocular eye rotation, to neurons representing monocular rotation. Notably, the anatomical left/right asymmetric distribution of the ROIs in the rotation clusters was consistent with previously described representation of ipsiversive movements in the hindbrain^18, 20, 25, 30, 42^.

## Discussion

Here we have studied hindbrain activity in zebrafish larvae performing a variety of eye and tail movements in response to a sequence of conjugate and disconjugate optokinetic stimuli. Using reduced rank regression, we found that the bulk of population activity in the hindbrain can be explained by two linear combinations of regressors, which we called latent regressors. Based on the contribution of these latent regressors to ROI activity, ROIs formed three main clusters. These clusters grouped ROIs whose activity represented vergent eye movement and swimming (1), and conjugate eye movement in clockwise (2) and counterclockwise (3) directions, as well as swimming. ROIs belonging to the three clusters showed a spatial distribution that was conserved across fish, suggesting that translation and rotational behaviors are encoded in separate circuits.

We used a stimulus protocol aimed at dissociating eye position and velocity, as well as left and right eyes. Indeed, using steps of constant velocity, the eye position space explored by the fish was expanded (Figure 1B), leading to both vergent and rotational eye movements (see also refs. 20,30), although divergent eye movements were less frequent. Moreover, the correlation between eye velocity and position was decreased (compared to the sinusoidally-modulated velocity stimulus), albeit not eliminated. The fish also generated forward and turning bouts in response to the stimuli (Figure 1D). Our stimulus protocol therefore succeeded in generating a large variety of behaviors. Among the 12 behavioral variables, some carried more variance than others, but the regressor space was still full-dimensional (Figures S4C,D).

In a previous study^30^, we showed that activity correlated with eye position, eye velocity and swimming was widely distributed across the brain. Several studies have reported neural activity related to these variables. Neurons tuned to (horizontal) eye position and velocity have been extensively studied in various motor nuclei, particularly in primates^54, 55^ and goldfish^56^, and more recently in zebrafish^18, 53, 57, 58^. The surveyed brain regions include the abducens and oculomotor nucleus, which directly innervate the muscles that move the eyes^13, 16, 17, 54, 55, 59^, as well as the oculomotor integrator, a brain area that transforms velocity commands into eye position^17, 53, 56, 57, 60–63^. Even though the majority of responses in these nuclei are monocular, there is a substantial fraction of neurons that represent eye movements binocularly^15, 18, 19^. The present study does not cover those brain regions, and direct comparison is therefore not possible. Velocity tuning during optokinetic behavior has been found in the goldfish and zebrafish hindbrain; whereas in adult goldfish a nucleus solely encoding eye velocity was found^64^, in zebrafish velocity encoding neurons show mixed selectivity, being also sensitive to position, and appear more distributed in the hindbrain (our results; see also ref. 18). Our findings suggest that encoding of eye velocity signals ranges from monocular to binocular (rotation) tuning.

The neurons we imaged similarly display a variety of tuning to eye position, velocity, and saccades, as well as tail movements, as shown by correlation maps (Figure 2; maps for individual regressors not shown, but see ref. 30). This apparent complexity at the single-neuron level essentially disappears when using reduced rank regression, which can be thought of as multivariate regression with built-in dimensionality reduction. Using reduced rank regression leverages the size of the data (thousands of neurons recorded over several hours) and identifies features that are common in the population, even if they are contaminated by irreducible imaging noise. Quite surprisingly, though we started with 12 regressors (then expanded to 732 through time shifts), neural activity in the hindbrain region we imaged was strongly dominated by only two modes of population activity, one related to vergence, and one to rotation. Moreover, these modes combined eye and tail movements. In turn, the diversity of tuning at the single-neuron level stems from different neurons expressing different combinations of the population modes (Figure S9), as well as imaging noise. Indeed, responses in our rotation clusters are diverse, and fall in a continuum going from monocular to conjugate (binocular) tuning of eye velocity. We believe that combining the measurement of many behavioral variables with this type of population analysis could also help to better understand the representation of eye and tail variables in some of the hindbrain areas we have not measured.

One caveat of our analysis is that by focusing on the dominant modes of population activity, we could squeeze out smaller, but still significant modes. For instance, even though the main modes capture over 90% of the explained variance, there could be smaller pools of neurons that are different and not captured by our analysis. While there were no strong outlier ROIs in our comparison of MLR and RRR (Figure 4B), there does appear to be more structure to the clusters in some of the fish (Figure S8). Future work will therefore have to see whether the clusters extracted here need to be refined.

We find a clear organization of the neurons into three different, anatomically separated groups. ROIs belonging to the rotation clusters were symmetrically distributed across the midline, and represented ipsiversive motion/position, as previously observed for eye and tail movements^22–25, 30^. The clusters mix sensory (stimulus velocity) and motor variables, suggesting that they are involved in sensorimotor transformations that segregate translational optic flow, vergent movement of the eyes, and forward swims, from rotational optic flow, conjugate eye movements, and (potentially) turning bouts. This is consistent with a role for the hindbrain in sensorimotor transformation, as previously postulated^22, 24, 32^. Interestingly, all three clusters mix tail and eye variables, suggesting that these neurons compute some higher-level motor commands that separate behaviors along one translational and two rotational axes.

Translational (vergent) and rotational motion could be directly fed to these populations, where they are combined with, or transformed into the corresponding behavioral output. Indeed, it has been shown that forward translational motion selectivity is already detectable in the pretectum^65–68^, and direct projections from the pretectum to premotor centers in the hindbrain^65^ could inform forward swimming. Although rotation selective neurons have been shown to be rare in the pretectum, a large fraction of neurons selective for leftwards/rightwards optic flow was found, suggesting that this area can also instruct left or right turns by projecting to a different set of premotor centers in the hindbrain. The same neurons that instruct left/right turns could mediate eye rotation. Notably, the subset of rotation-selective neurons we observe could receive direct input from the (few) rotation-selective cells in the pretectum^65–68^, or their binocular tuning could result from combining the more dominant monocular pretectal signals.

That neuronal activity in this area can be assigned to two different principal modes, one related to rotational (conjugate) motion and one related to translation (vergent) motion, is consistent with a binocular organization of the control of eye movements, as postulated by Hering^9, 10^, at least in the areas imaged here. However, it is important to note that in the rotational clusters responses showed a gradient from binocular to monocular. The monocular responses observed in downstream areas (oculomotor integrator, abducens nucleus) could reflect a combination of conjugate and vergence signals provided by the neurons imaged here, or could be directly informed by the subpopulation displaying monocular responses.

One potential caveat here is that the regression method may have failed to separate the (highly correlated) stimulus and eye velocities. Theoretically, regularized multiple linear regression will separate correlated variables (as long as they are not perfectly correlated). However, if the relation between regressors and behavioral variables is non-linear, then such a separation is not guaranteed. Moreover, the eye velocity regressor, which is estimated from noisy eye positions, is less reliable than the stimulus velocity regressor, such that its overall importance for ROI activity may have been underestimated. In turn, it is possible that the vergence cluster is, in truth, really related to retinal slip, whereas the rotation cluster is only related to eye velocity. More targeted experiments will be able to resolve that question.

In summary, we show here that using reduced-rank regression, which combines dimensionality reduction with multilinear regression analysis, we could leverage large data sets to reveal the dominant modes of behavior-related activity present in the hindbrain population we imaged. We found that the bulk of hindbrain activity when responding to different patterns of whole-field rotation was distributed in three populations of neurons, representing translational and left/right rotational information. This method can be useful to reveal the main modes of population activity and reduce the complexity of datasets in large scale recordings in behaving animals.

## Acknowledgments

We thank João Semedo for helpful insights in the RRR analysis. Owen Randlett for providing the Z-brain rhombomere masks. José Lima for the initial version of the tracking software. Lucas Martins for help with the visual stimulation and tracking software. Misha Ahrens for providing the *elavl3*:H2B-GCaMP6s line. Ruth Diez del Corral, Adrien Jouary and Gokul Rajan for comments on the manuscript. We thank the Champalimaud Research Fish, Software and Hardware platforms for logistical support, and the Communication, Events and Outreach team for help with schematics. This work was realized through funding to MBO from an ERC Consolidator Grant (Neurofish) and the Portuguese Fundação para a Ciência e Tecnologia (FCT) (PTDC/NEU-SCC/5221/2014), to CEF, MBO, and CKM from the Champalimaud Foundation, to RP and MBO from the Volkswagen Stiftung Life? Initiative and to RP from the DFG under Germany’s Excellence Strategy within the framework of the Munich Cluster for Systems Neurology (EXC 2145 SyNergy -- ID 390857198). We received support from the research infrastructure CONGENTO, co-financed by Lisboa Regional Operational Programme (Lisboa2020), under the PORTUGAL 2020 Partnership Agreement, through the European Regional Development Fund (ERDF) and Fundação para a Ciência e Tecnologia (Portugal) under the project LISBOA-01-0145-FEDER-022170.

## Author contributions

C.E.F., M.B.O. and R.P. conceived the experiments. C.E.F., M.dG. and R.P. performed the experiments. C.E.F. and C.K.M. conceived the analysis. C.E.F analyzed the data. A.L. developed the behavioral tracking software. A.O. performed brain registrations. C.E.F. and C.K.M. wrote the manuscript. All authors provided feedback on the manuscript.

## Methods

### Fish care

Adult fish were maintained at 25⁰C on a 14:10 hour light cycle following standard methods^69^. Embryos were collected and larvae were raised at 28⁰C in E3 embryo medium (5 mM NaCl, 0.17 mM KCl, 0.33 mM CaCl2 and 0.33 mM MgSO4) in groups of 30. Transgenic lines: *elavl3*:H2B-GCaMP6s (Vladimirov et al. 2014), in a nacre (*mitfa* -/-) background were used. All experimental procedures were approved by the Champalimaud Foundation Ethics Committee and the Portuguese Direcção Geral Veterinária, and were performed according to the European Directive 2010/63/EU.

### Calcium imaging and behavior recording

Six and seven day-old zebrafish larvae were placed in a drop of 1.6% low-melting-temperature agarose (1.6% UltraPureTM LMP Agarose, Invitrogen 16520-100, in E3 water), in a Petri dish with a Sylgard 184 base (Dow Corning) and immersed in E3 water. The agarose around the tail, caudal to the pectoral fins, and eyes was cut away with a fine scalpel to allow for eye and tail movement. The dish was placed onto a light-diffusing screen and imaged on a custom-built two-photon microscope. A Ti-Sapphire laser (Chameleon, Coherent) tuned to 950 nm was used for excitation; power at the sample ranged between 4.2 and 5.6 mW. Frames were acquired at 2 Hz. Each plane was imagined for approximately 125 seconds, the time required for the complete stimulus set (see below). Pixel size was 0.53 µm for x and y. The imaged plane was then shifted by 1 µm.

### Visual stimulation

Visual stimuli were displayed using a custom-written rendering engine using OpenTK and the stimuli were generated using OpenGL Shaders (Alexandre Laborde and Lucas Martins, unpublished), and projected at 60 frames per second using an LED projector (Optoma Europe Ltd.). The output of the projector was filtered to allow for simultaneous imaging and visual stimulation (long-pass colored glass filter Thorlabs FGL590 and a TXRED emission filter Thorlabs MF630.9).

The stimulus consisted of radial red and black stripes with a period of 45 degrees. The stimulus was centered on the fish, and was virtually split in two, so that each eye could be stimulated independently. To ensure that the stimulus in one hemifield could not be seen by the other eye, an area in front of the fish spanning an angle of 57.3 degrees was kept unstimulated (black; see schematic in Figure 1A and Movie S1). The rotation of the stimulus in each screen-half was modulated independently, with steps of constant velocity (-30, -15, 0, 15, 30 °/s) lasting 5 seconds. A stimulus set or repetition, which consisted of the 25 possible combinations of velocities, was presented in each imaging plane. For each repetition, the order of the velocity pairs varied randomly.

### Behavior tracking

To track eye and tail movements, a small hole was cut in the diffusing screen and the fish was imaged from below using a high-speed, infrared-sensitive CCD camera (Mikrotron EoSens® CL 1362) coupled to a 50 mm objective lens (Schneider Kreuznach Xenoplan 2.8/50-0902) and a pair of filters: a 800 nm short-pass filter (Edmund Optics 64333) and a long pass-filter LP695 (Edmund Optics 32756). Fish were illuminated from above using an infrared LED (M780L2, Thorlabs) through an aspheric condenser diffusing lens (Thorlabs ACL5040-DG6-B). Additional illumination for the tail was provided by two infrared LEDs (780 and 800 nm) positioned close to the tail.

Tail data was acquired at 700 Hz, while eye data was acquired at 100Hz. Tail and eye position were extracted online using custom written software (C#). Briefly, eye centers were marked before the beginning of the experiment. The eye object was extracted based on the pixel intensity using the flood fill algorithm. After that, the orientation of the eye object was approximated by calculating the first and second order central image moments. The eye angle was obtained by the angle of the major axis of the approximated eye object relative to the horizontal plane of the image^70^. The angles were then corrected to be defined relative to the midline of the fish; temporal positions were defined as positive (Figure S1A). The starting point of the tail was marked at the caudal part of the swim bladder, and the endpoint was the tip of the tail. The tracking software created 16 points in between. Tail angles and position, as well as eye angles, were saved to text files for offline analysis.

### Behavior analysis

All data analysis was performed in MATLAB (MathWorks, USA).

Bouts of swimming were extracted from tail angle data as follows. The angle of the tail relative to the body was obtained by calculating the cumulative sum of the angle differences of the first 12 segments (the last 4 segments were excluded as they were too noisy). We then calculated tail vigor as a rolling standard deviation in a window of ∼44 ms (30 frames) of this trace (after ref. 71). Bouts were defined as the segments when tail vigor was above a threshold; this threshold was chosen for each fish, based on the distribution of tail vigor values.

To calculate the tail turning bias (simply called bias), tail traces were centered by subtracting the median tail deflection angle. Bias was then calculated for each bout. For a bout, bias was defined as the sum of each tail segment deflection angle, divided by the absolute value of that sum (after ref. 32). Bouts were classified as forward swims, or left or right turns according to their bias value and a given threshold; thresholds were chosen manually for each fish by analyzing the mean tail traces associated with different threshold values. Absolute bias values greater than the threshold were classified as right/left turns, whereas bias values in between were classified as forward swims. Positive values of bias corresponded to a net rightwards tail bent, while negative values corresponded to a leftwards tail bent.

Eye position was defined as the angle relative to the midline of the fish. Eye position traces were first filtered using a smoothing polynomial filter (Savitzky-Golay filter, order 5, width 15) to denoise the data while preserving the saccades. The data was further smoothed using a medial filter in a small window (medfilt1, width 5, corresponding to 50 ms), to further reduce the noise.

Eye velocity was estimated using a differentiator filter to avoid the introduction of high frequency noise when differentiating. Eye position and velocity were then interpolated to match the rate of tail acquisition (700Hz), so that all pieces of behavioral data could be temporally aligned.

Eye velocity was decomposed into low, ‘following’ velocity, and high velocity, which corresponded to saccades. To do so, the velocity trace was first filtered using a median filter (medfilt1, in a window of 9 frames, corresponding to approximately 15 ms), and saccades were defined as points in which velocity was above a threshold, chosen individually for each fish (typically, thresholds were 2 to 4 standard deviations above the mean velocity value). A ‘high velocity’ or saccade trace was then generated by keeping only the velocities above threshold, and setting the rest of the trace to 0. A low velocity trace was generated by removing values above threshold from the velocity trace, and replacing them using the ‘fillmissing’ (‘nearest’) function in MATLAB. The low velocity trace was further filtered using a running average of 150 points (approximately 250 ms). Temporal eye positions and velocities were defined as positive for both eyes (Figure 1A, S1A).

Eye gain was defined as the ratio of eye velocity divided by the corresponding stimulus velocity. Vergence index was defined as the ratio of the variance of vergent eye positions over the variance of rotation eye positions.

We selected 12 behavioral variables. The variables included were left/right eye position, left/right (low) eye velocity, left/right high eye velocity (saccades) associated with a bout, left/right high eye velocity (saccades) not associated with a bout, left/right stimulus velocity, tail vigor (which is a measure of swimming) and tail bias (which is a measure of laterality of the tail flick).

### Image analysis

Image analysis was performed with MATLAB (MathWorks, USA). Images were corrected for drift or small movements of the fish as previously described^30^. Briefly, each image frame was aligned, using translation only, to the average image of that *z*-plane. Because an interlaced scanning was used, images were aligned first in the horizontal dimension only. Vertical alignments were subsequently performed, but restricted to an integer number of lines. Information about the vertical displacements applied could be used later for reconstructing temporal data. After aligning all frames of a given plane to the average of that plane, consecutive *z*-planes were aligned to each other with subpixel precision. Occasionally vigorous swimming movements caused a large motion in a single frame. These frames were detected based on (decreased) correlations between consecutive frames, and replaced with a linearly-interpolated frame (using the fillmissing function in MATLAB).

Stacks were segmented in overlapping voxels (that we will call ROIs) in the form of cuboids, of 9 x 9 x 9 pixels (4.77 µm x 4.77 µm x 9 µm), tiling the brain in steps of 3 (x,y) or 3 (z) voxels. Because each voxel contained 9 planes, dF/F was computed first for each plane, with F the mean across frames and pixels in a given plane, and dF the difference between the fluorescence value of a given pixel and F. When pulling together *z* planes in a given voxel, dF/F was centered by subtracting the mean across the planes included. This centered dF/F value was used for the subsequent analysis.

Planes that showed motion artifacts that could not be corrected were excluded from the analysis. If a voxel contained one such plane, the part of the trace corresponding to that plane was excluded.

### Correlation analysis

Using the initial 12 behavioral variables, we first generated an expanded set of 155 regressors by thresholding eye positions and velocities with various thresholds, by computing eye acceleration, by combining left and right variables, etc. (see Table 1). These variables were downsampled to allow for straightforward alignment to imaging data.

For correlation analysis (Figure 2E-F), we accounted for delays introduced by the calcium indicator dynamics by first convolving regressors with a kernel with an exponential decay based on the measured half-decay time for nuclear GCaMP6s (4.68s; our observations), to produce a set of predicted fluorescence traces^30, 42^. These traces were then compared with the measured fluorescence traces by correlation. Systematic correlation analysis was performed on cuboidal ROIs of 9 x 9 x 9 pixels.

### Regression analysis

#### Criterion for ROI selection

ROIs were pre-selected based on the correlations with the expanded (155) set of regressors: ROIs that had a correlation greater than a given threshold with any of the regressors were included in the regression analysis. A threshold of 0.4 was chosen for the analysis in this paper. This resulted in an average of 43000 (ranging from 27000 to 57000) ROIs in each fish. Strictly speaking, a preselection is not necessary for the analysis, as noisy or uncorrelated ROIs do not contribute to the estimation of the features anyways. However, we performed the preselection to maintain computational feasibility, and we checked that manipulating the selection threshold did not affect the results. As a control, we ran the analysis both on a randomly-selected subset of ROIs, and using different thresholds for preselection. In both cases, the results were not significantly different from those using ROIs selected based on correlations (not shown).

#### Regressors

Regressors related to left/right variables were expressed as vergence (left+right) and rotation (left-right); i.e., instead of using ‘left eye position’ and ‘right eye position’ as regressors, we used ‘eye position vergence’, which is simply the sum of the left and right eye positions, and ‘eye position rotation’, which is the difference between left and right eye position (Figure S2). This represents a 90 degree rotation of the axes (white axes in Figure 1B; Figure S1A).

For regression analysis, regressors were ***not*** convolved with the indicator kernel. Rather, to account for the delay introduced by the indicator, regressors were shifted 20 seconds into the past and 10 seconds into the future, in steps of 0.5 seconds, creating an expanded set of 61 regressors (Figure 3) for each of the 12 regressors. Altogether, we therefore used 732 linearly-independent regressors. Singular value decomposition of the z-scored regressors shows that more than 12 components of the regressor matrix have significant power (Figures S4C-D).

Regressors were normalized (z-scored) using the mean and standard deviation of the training set (see *Estimation of hyperparameters* below). For each voxel, fluorescence traces were centered by subtracting the mean fluorescence of its corresponding training set.

#### Reduced rank regression (RRR)

We sought to relate the measured activity to our set of regressors using multiple linear regression (Figure 3 and Figure S3). To reduce the dimensionality of the population data, we then asked whether activity could be explained by a subspace of our regressor space. To answer this question, we used reduced-rank regression^39, 48, 49^ (RRR).

We started with standard multiple linear regression (MLR), which relates the M×T matrix of regressors (M regressors, T timepoints), *X*, to the N×T matrix of fluorescence traces (N ROIs, T timepoints), *Y*,

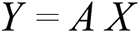

where *A* is a N×M matrix of regression coefficients (Figure S3A). *A* can be found using the ordinary least squares (OLS) solution so that

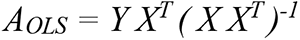

To reduce overfitting, we use Ridge Regression^51^, which introduces a regularization term in the matrix inverse so that now

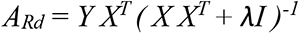

where I is a M×M identity matrix and *λ* is a constant that establishes the degree of regularization. The predicted ROI traces (or predictors) are then given by

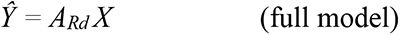

Because in our experiments planes were recorded sequentially, the matrix *Y* is incomplete (i.e., we have values for the regressors throughout the experiment, but for each ROI, activity was only measured during part of the experiment). Doing the OLS regression on a matrix of N dependent variables is equivalent to doing N separate OLS regressions, one for each of the dependent variables. Therefore, we do a Ridge Regression for each ROI (using its corresponding regressor trace), and obtain a set of regression coefficients for that ROI. We then construct the matrix *A_Rd_* by placing each set of coefficients in a row. This matrix *A_Rd_* is now used to calculate *Ŷ*. To ensure statistical comparability across ROIs measured in different planes, both the regressors and the activity should ideally be stationary. To ensure stationarity of the regressors, we first analyzed the covariance matrix of the regressors across pairs of planes (Figure S4A,B). We then asked whether the regressors for a given plane were ‘different’ from those of the other planes by looking at the distance of the covariance matrix of that plane to those of the rest (Figure S4A). Planes were considered different from the rest if the mean distance to all other planes was above a threshold (defined as f×standard deviations above the mean; Figure S4B); those planes were removed from the analysis. The stationarity of ROI activities across planes could not be tested, as we did not measure activity of individual ROIs throughout the whole experiment.

To find a subspace of the regressors that could be sufficient to predict *Y*, we used reduced-rank regression (RRR), which seeks to minimize the mean-square error under a rank constraint on the matrix of regression coefficients, *A*. Denoting the rank of such a matrix with the index *q*, we now seek to solve:

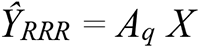

such that *Ŷ_RRR_ is close to Y in the mean-square sense.* This can be solved by calculating the singular value decomposition of the predictors *Ŷ* obtained using Ridge-regularized OLS as above,

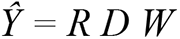

And then setting

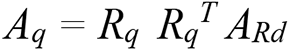

where *R_q_* is a matrix composed of the first *q* columns of the *R* matrix from the singular value decomposition of *Ŷ*.

Now we can predict fluorescence activity using:

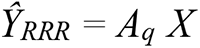

The bases of the subspace of ‘latent’ regressors or *features* that come up in our model is given by a singular value decomposition of the matrix of regression coefficients, *A_q_*:

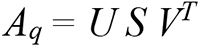

We can therefore rewrite the predicted ROI activity as

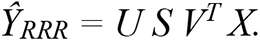

Where *U* has size N×Q and *V^T^* has size Q×M. In turn, we can interpret

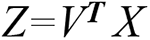

as a ‘new’ and much smaller set of *q* regressors that can be used instead of the M original regressors, *X*. This new set of *latent* regressors is built from a linear combination of the original regressors *X*, with weights given by the columns of the matrix *V* (Figure 3B,C). We will call these columns the *features* associated with the latent regressors. Scaling the features *V* by multiplying them with the corresponding singular value S results in a scaled version of the feature, that highlights the relative importance of the feature, and we use that form in Figures 4 and S6. The columns of the matrix *U* indicate how much each of the latent regressors contributes to the activity of each ROI (Figure 3B; Figure S3B). We call these columns the *(feature) contributions*.

##### Estimation of hyperparameters

Rank *q* (the number of features) and the regularization parameter *λ* were determined using five-fold cross-validation. For each round of cross-validation, 80% of the data was used as a training set, while the remaining 20% of the data was used as a test set. Data was taken in blocks and not randomly sampled to preserve the temporal structure of the traces. In each iteration, we selected the smallest rank (number of features) and the largest *λ* for which the performance of the model (see below for definition) was within one standard error of the maximum performance (‘one standard error rule’ ^51^; Figure S5A).

Model performance or explained variance (EV) was computed for each ROI as

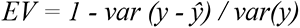

where *y* is the 1×T fluorescence trace for a given ROI and *ŷ* is the 1×T predicted fluorescence trace for that ROI using the full (Ridge-MLR) or the reduced (Ridge-RRR) model.

The average model performance or population explained variance (popEV) was computed as

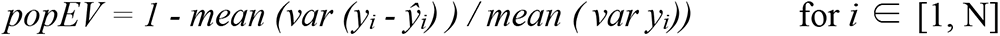

where *y_i_* and *ŷ_i_* are the traces for a given ROI *i*, and the mean is that across all n ROIs. popEV was used to select the model parameters *q* and *λ*.

We built separate RRR models for each fish (8 fish). The inputs to the model were a matrix of 732 regressors (*X*) and a matrix of fluorescence traces (*Y*) of an average of 43000 ROIs (see above for selection criteria and RRR formulas).

As a control, and to assess the significance of the contribution (*U*) values in our models, we built a model where regressors were temporally shuffled to disrupt the relationship with ROI activity. Regressors were shuffled by randomly sorting blocks of 100 points. The resulting model could not explain the data (Figure S5B), and the latent regressor obtained had on average a contribution of zero to population activity (compare *U* values for the model with real data and that using shuffled regressors, for the same fish; Figure S5C).

### Clustering

ROIs were clustered in two ways. First, we clustered ROIs manually. Clustering was performed in feature contribution (*U*) space, in scatter plots for the contributions of two of the first three latent regressors (we chose the latent regressors that led to the clearest clustering of ROIs; in most cases, this corresponded to the latent regressors associated with features 1 and 2 (6 out 8 fish; Figure 5A); in two fish this corresponded to the latent regressors associated with features 1 and 3. Clusters were defined by manually drawing polygons around the data points (Figure 5B-C), following the valleys in the density maps. Only ROIs for which the model’s explanatory power was greater than 0.4 were included in the clustering.

Second, ROIs were clustered using ClusterDv^52^, a density-based clustering algorithm that can automatically find the best number of clusters. Clustering was done in 2, 3, 4 and 5 dimensions (when existing), yielding qualitatively identical results. The results using ClusterDv and manual clustering were indistinguishable (not shown).

### Anatomical registration

#### Registration

All registrations were run on Ubuntu server 18.04.4 as a Windows Subsystem Linux instance, using the Computational Morphometry Toolkit (CMTK – www.nitrc.org/projects/cmtk/) version 3.3.1. The host PC was using a 6 core AMD Ryzen 5 2600 Processor with 64Gb of RAM.

#### Template

Whole brain anatomical stacks of *elavl3*:H2B-GCaMP6s transgenic zebrafish were obtained by averaging all frames corresponding to a plane. These stacks were acquired with a voxel spacing of 1.13μ x 1.13μ x 1.0μ, and include almost the entire brain (with the exception of the most rostral part of the telencephalon). Seventeen larvae were imaged (6 d.p.f. and 7 d.p.f.), and the seven most complete stacks were selected for template generation.

Templates were generated as follows: one anatomical stack was chosen as a seed, and all other images were registered to this seed using CMTK’s registration and warp functions using CMTK’s provided wrapper script — munger.pl --- (adapted from code at https://github.com/jefferislab/MakeAverageBrain; Greg Jefferis) with options “-X 26 -C 8 -G 120 -R 3”. Additional options passed to the Affine and Warp functions were -A “--sampling 2 -- accuracy 4 --omit-original-data” and -W “--sampling 2 --accuracy 1.5 --omit-original-data”, respectively. When all of the registrations were complete, they were used as input to CMTK’s avg_adm function to create a shape-averaged greyscale average. This average was then used as the seed for another round of registrations. This process was repeated a total of three times, and the output of the last shape-averaging step was used as the whole-brain *elavl3*:H2B-GCaMP6s template in subsequent steps.

#### Registrations of functional imaging data to the anatomical template

Hindbrain functional data was registered to the *elavl3*:H2B-GCaMP6s whole-brain template. For each fish, a hindbrain anatomical stack was generated by averaging all frames of a given plane. Hindbrain images had a voxel spacing of 0.53μ x 0.53μ x 1.0μ. After some experimentation, it was determined that these stacks could be registered directly to the anatomical whole brain template by passing a cropping argument (’--crop-index-ref’) to CMTK’s Registration and Warp functions.

Registrations were performed using munger.pl with options “-X 26 -C 8 -G 120 -R 3”. Additional options passed to the Affine and Warp functions were -A “--sampling 2 --accuracy 4 --crop-index-ref 33,89,2,380,391,195” and -W “--sampling 2 --accuracy 1.5 --crop-index-ref 33,89,2,380,391,195”, respectively. Finally, to account for excess warping in some of the stacks after registration, munger.pl’s “-E” switch was used with some of the registrations to further constrain the weight for grid bending energy. This parameter can vary between 0 and 1, and in these cases was varied from the 0.1 default, up to 0.75. Finally, registrations where the bending energy did not equal 0.1 also used the default ‘sampling’ and ‘accuracy’ for the affine and warp steps. Registrations that had good overlap with the template and minimal warping were used in subsequent steps.

The best registrations derived from the above process were then used to drive transformations of the hindbrain anatomical and analysis (correlation, features) stacks associated with that larvae/experiment. Specifically, a shell script (reformat_loop.sh) organized the images and the warp output files in order to run CMTK function ’reformatx’. Correlation and feature maps were registered to the hindbrain template using CMTK’s reformatx function, and nearest neighbour (— nn) interpolation was used in order to preserve the values of the voxels as they were mapped into the reference space

#### Bridging registrations

The whole-brain *elavl3*:H2B-GCaMP6s template was registered to the ZBB ^72^, ZBrain ^47^, and MPI (https://fishatlas.neuro.mpg.de/) templates using CMTK. To account for differences in the rotations and *z* orientations of the whole-brain *elavl3*:H2B-GCaMP6s template to the external templates, the CMTK registration was seeded with rigid transformations that describe the necessary rotating and flipping of the images. For example, see script munger_bridge_f4wb_mpin.sh; this script seeds the affine transformation with a rigid transformation that rotates the brain to match the rostral/caudal orientation of the MPI template, as well as inverting the z-axis to account for the difference in the order of acquisition of the stack. To account for some undesired warping during bridging to the MPI(2) template, the bridging registration was updated to use an energy constraint of 0.5, thus improving the reformatting of individual hindbrains to the MPI(2) template.

