## Supplemental Data for "Dimensionality reduction reveals separate translation and rotation populations in the zebrafish hindbrain"

**Supplementary Material**

March 2023

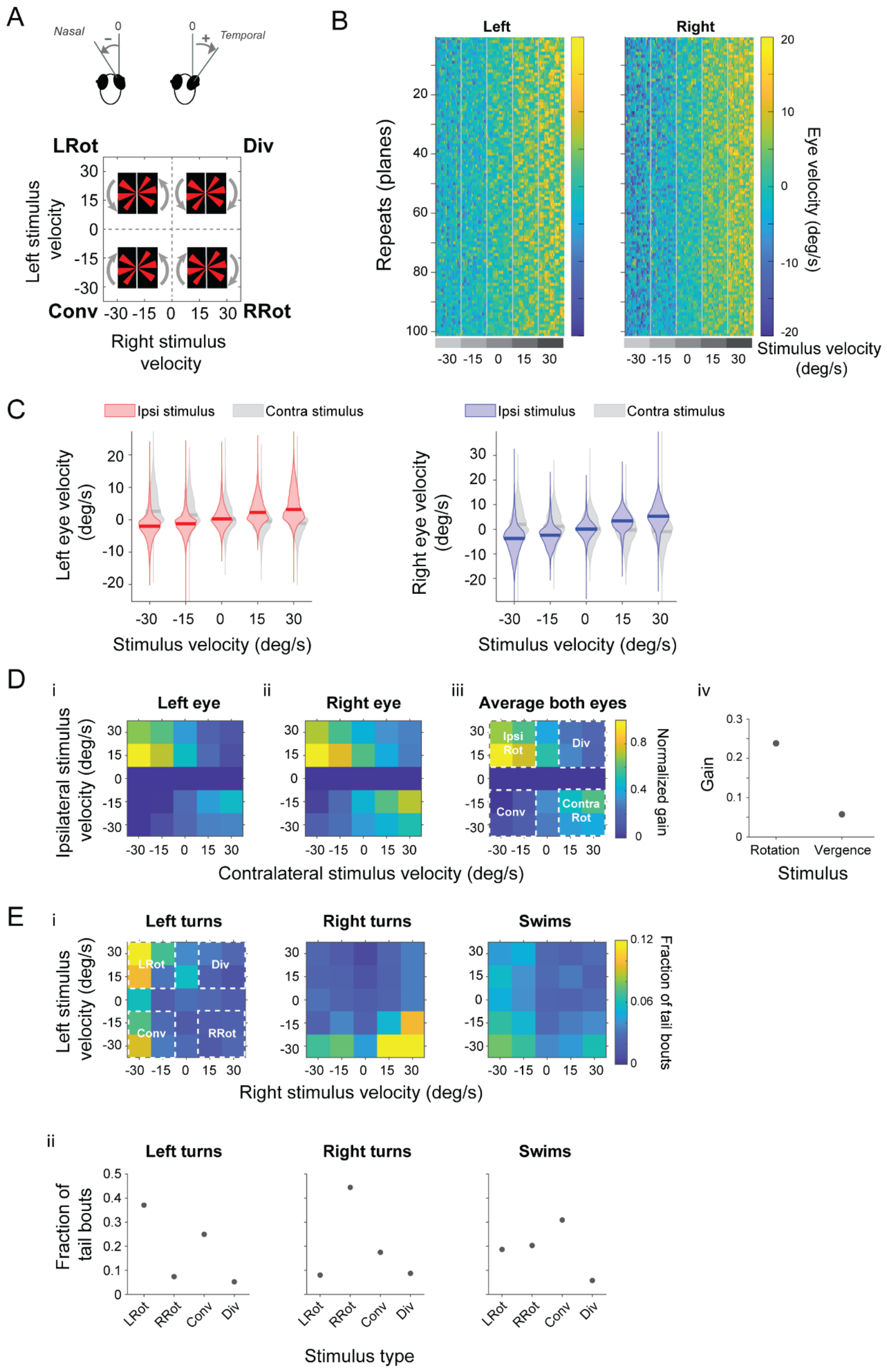
A
Nasal
Temporal
LRot
Div
Conv
RRot
Left stimulus velocity
Right stimulus velocity
B
Left
Right
Repeats (planes)
Eye velocity (deg/s)
Stimulus velocity (deg/s)
C
Ipsi stimulus
Contra stimulus
Left eye velocity (deg/s)
Right eye velocity (deg/s)
Stimulus velocity (deg/s)
D
i
Left eye
ii
Right eye
iii
Average both eyes
Ipsi Rot
Div
Conv
Contra Rot
Normalized gain
iv
Gain
Rotation
Vergence
Stimulus
Ipsilateral stimulus velocity (deg/s)
Contralateral stimulus velocity (deg/s)
E
i
Left turns
Right turns
Swims
LRot
Div
Conv
RRot
Fraction of tail bouts
Left stimulus velocity (deg/s)
Right stimulus velocity (deg/s)
ii
Left turns
Right turns
Swims
Fraction of tail bouts
LRot
RRot
Conv
Div
Stimulus type

**Figure S1**. **Eye velocity and swimming are modulated by the stimulus.** Behavior for one example fish. **(A)** Top, Schematic showing the sign convention used: Temporal (T) positions/velocities are positive, nasal (N) positions/velocities are negative. Bottom, Schematic showing the different stimulus categories, according to the sign of the velocities presented in both hemifields: LRot, leftwards rotation; RRot: rightwards rotation; Conv: converging; Div: diverging. **(B)** Eye velocity (after removing saccades, 'low velocity') for the left and right eyes (left and right panels, respectively), sorted according to the velocity of the stimulus presented to that eye. Each row corresponds to a repetition of the stimulus set (note that in the experiment stimulus order was randomized in each repeat; here they are sorted to emphasize the modulation of behavior). **(C)** Distribution of low eye velocity for left and right eye (left and right panels, respectively), as a function of the velocity of the ipsilateral stimulus (red and blue), and the contralateral stimulus (gray). Dark lines indicate the median velocity for the stimulus condition. **(D)** Velocity gain as a function of ipsilateral and contralateral stimuli, for the left (i) and right eyes (ii), and for the average of both eyes. (iii) Each 2D-plot is divided in four quadrants corresponding to the four stimulus categories (dotted squares): ipsiversive rotation (IpsiRot), contraversive rotation (ContraRot), convergence (Conv) and divergence (Div). (iv) Average gain for rotating and vergent stimuli, for both eyes pooled. **(E)** Swimming bout type is modulated by stimulus category. (i) Frequency of left and right turns, and forward swims (see Methods), as a function of both left and right stimulus velocity, for the same example fish as in (D). (ii) Each 2D-plot is divided in four quadrants corresponding to the four stimulus categories (dotted squares): leftwards rotation (LRot), rightwards rotation (RRot), convergence (Conv) and divergence (Div), and the bouts inside each category are pooled.

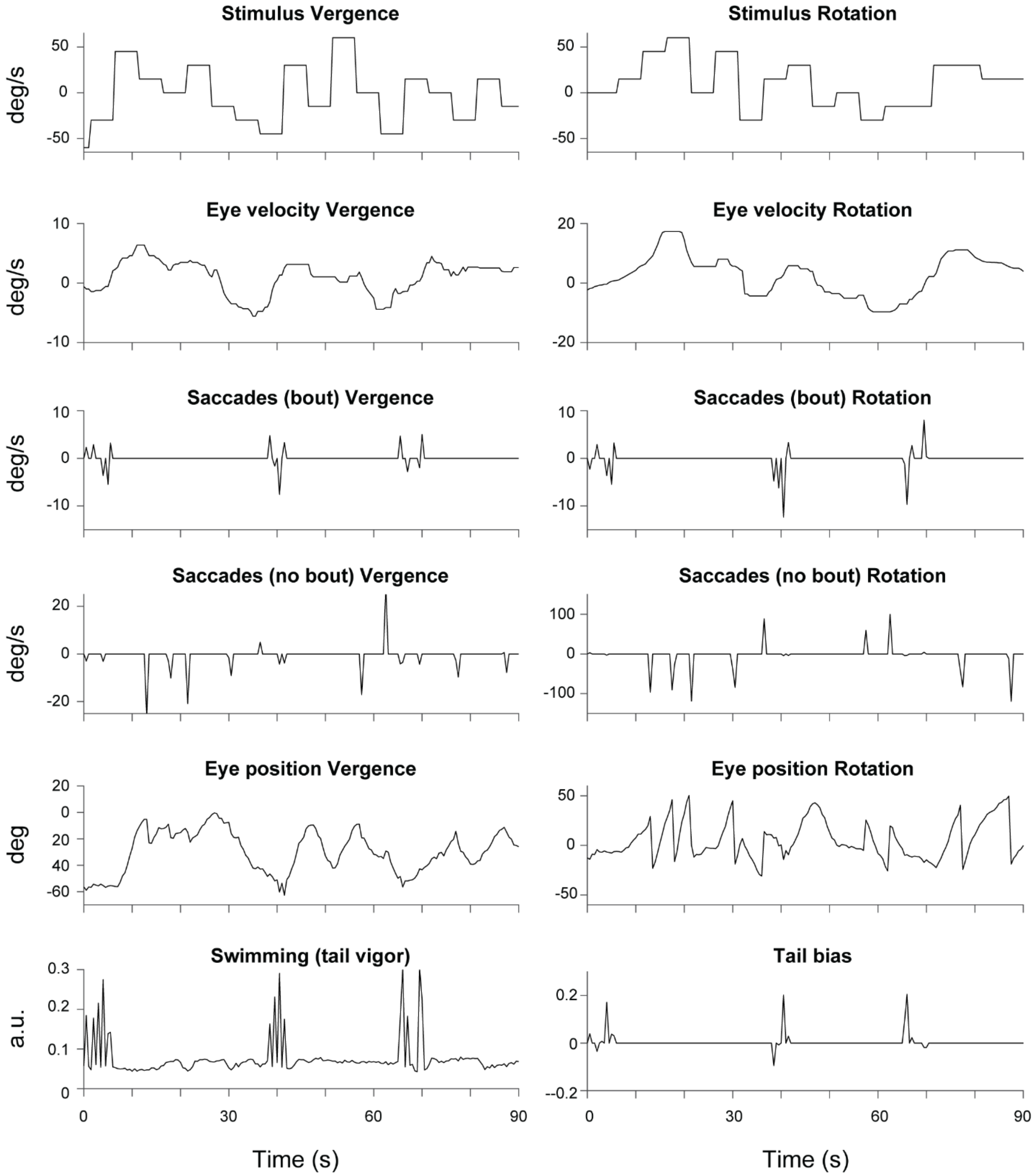


**Figure S2. Variables used as regressors for MLR/RRR analysis.** Eye position and tail angles were extracted online during the experiment, and used to derive the variables used for analysis (see Methods). Short representative traces for an example fish are shown.

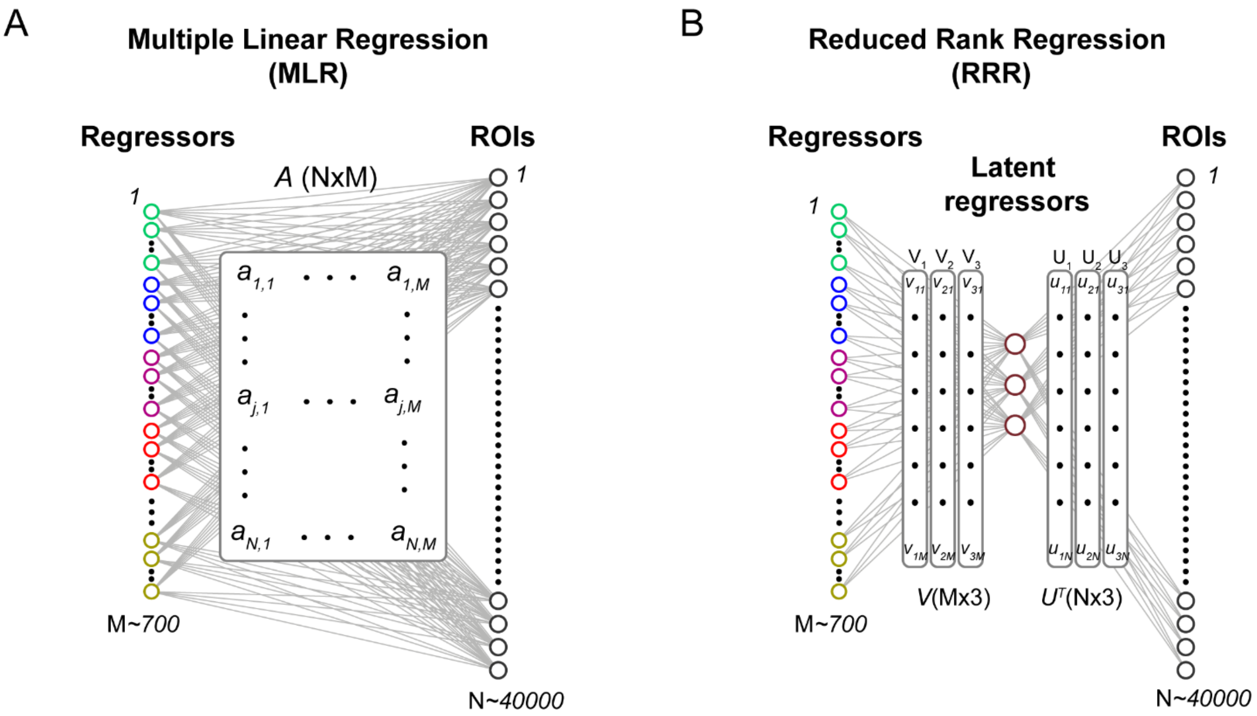


**Figure S3. Comparison of multiple linear regression (MLR) and reduced-rank regression (RRR). (A)** For a MLR model with M regressors and N ROIs, the model estimates a coefficient matrix *A* of size N×M; for each ROI, the corresponding row of *A* contains the weights for each regressor and its time shifts. **(B)** Using reduced-rank regression (RRR; see Methods) for the ROI population, we find a mapping from the M~700 regressors onto a small set of new regressors (rank 3 in this schematic; rank 3 to 6 in our data), such that these new regressors can explain the activity of all ROIs. Each of these new regressors is a linear combination of the regressors, with weights given by the features *V*. For each ROI, we can then determine the contribution *U* of each new regressor.

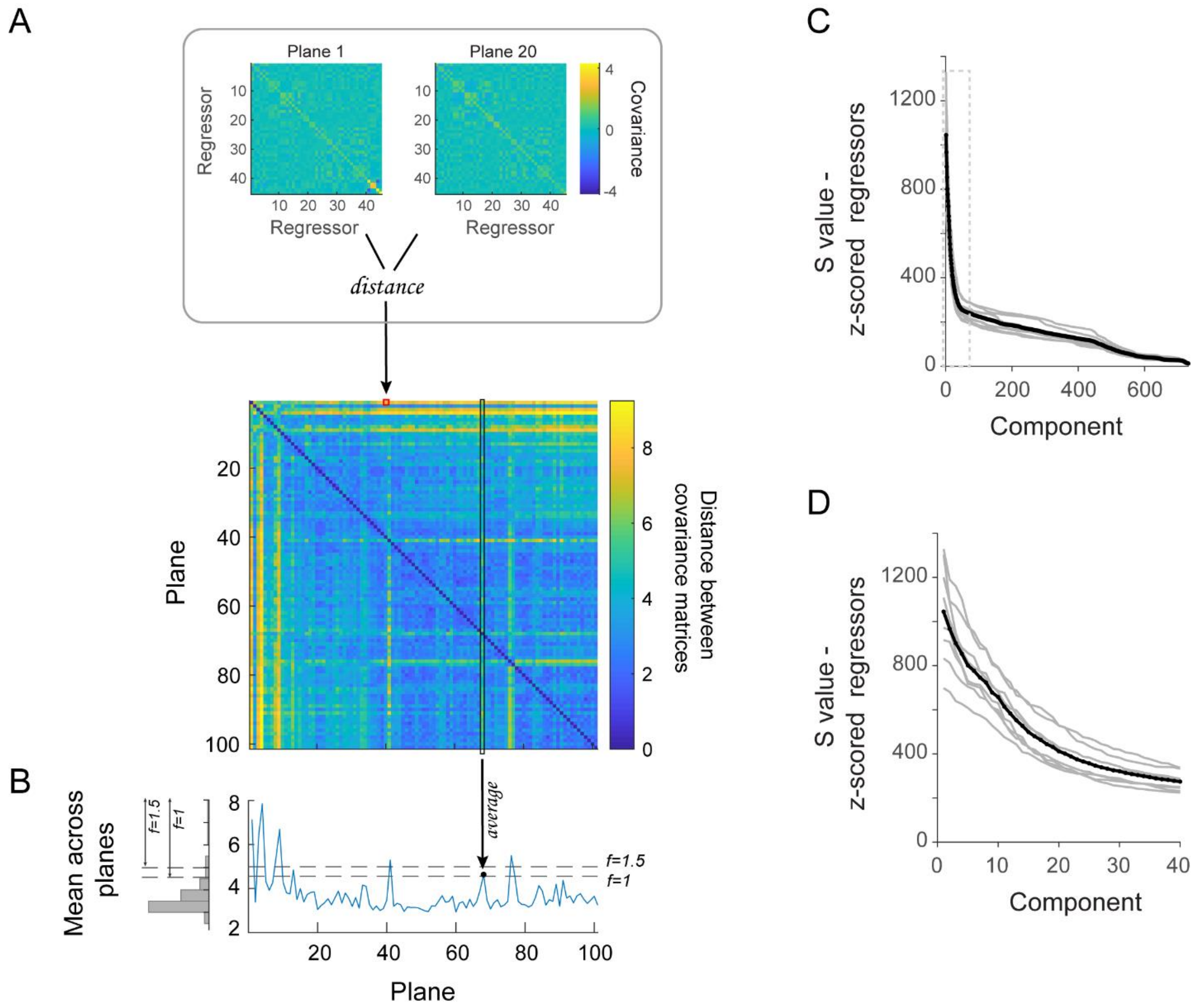
A
Plane 1
Plane 20
Regressor
Covariance
distance
Plane
Distance between covariance matrices
B
Mean across planes
average
f=1.5
f=1
C
S value - z-scored regressors
Component
D
S value - z-scored regressors
Component

**Figure S4. Regressors properties. (A-B)** Stationarity of regressors. In order to stitch together different imaging planes for the regression analysis, we needed to ensure that behavior was stable across planes. The stability of behavior was assessed using the stationarity of the regressors. We excluded from the analysis those planes for which regressors were 'different', as shown here. (A) Distance between the covariance matrices of regressors for each pair of planes. Inset illustrates how each element of the matrix is calculated. (B) Distance averaged across planes (black box) for each plane. Planes for which this mean was higher than a threshold (mean+f×std) were excluded from analysis. **(C-D)** The regressor space is multidimensional. Singular value decomposition (SVD) of the regressor matrix shows that the dimensionality of the regressor space is higher than 12 (the number of original regressors). (C) Singular value for the different components for SVD performed on the matrix of raw (unnormalized) regressors. (D) Same as in (C) but zoomed in for the first 40 components. Note that the regressors span several dimensions: the elbow of the SVD plots occurs around 15-20 dimensions. Gray traces, individual fish (N=8); black trace, mean across fish.

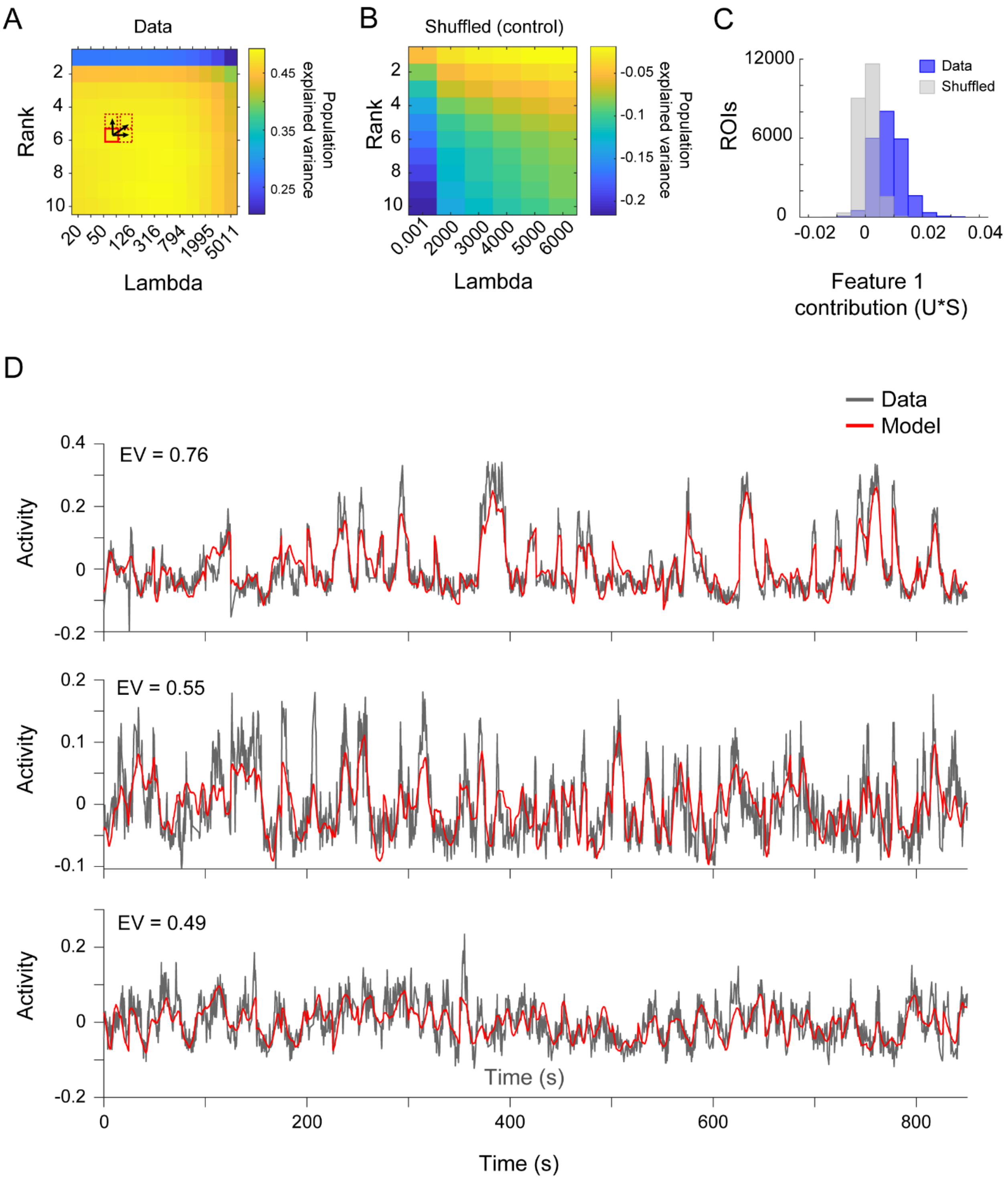

A
Data
Rank
Lambda
Population explained variance
0.45
0.35
0.25
B
Shuffled (control)
Rank
Lambda
Population explained variance
-0.05
-0.1
-0.15
-0.2
C
Data
Shuffled
ROIs
Feature 1 contribution (U*S)
D
Data
Model
EV = 0.76
EV = 0.55
EV = 0.49
Activity
Time (s)

**Figure S5. Model selection and performance. (A)** The regularization parameter **λ** and the regression matrix rank *q* were selected to maximize population explained variance (popEV, see Methods) in the test set, as follows: For each combination of **λ** and rank, the mean popEV in the test set (color scale) across five cross-validation sets is calculated. To select rank and **λ**, we first find the **λ**/rank combination with the highest popEV (red rectangle). We then choose the lowest rank and highest **λ**s for which the explained variance is within one standard error of the maximum (dotted areas; 'one standard error' rule, Hastie, Trevor et al., 2009). In this example, we select the model with rank 6 (six features) and **λ**=2000. **(B-C)** Shuffle control. Abolishing the relationship between regressors and activity yields a model with no explanatory power. Data from the same fish as in (A), but where regressors were temporally shuffled (see Methods). (B) Population explained variance of the shuffle control as a function of rank and lambda. (C) The contribution of the first (and only) new regressor in the shuffle control averages to zero (gray); compare with the distribution of feature 1 contributions for the model using real data (blue; for the same set of ROIs in the same example fish). **(D)** Comparison of real (gray) and predicted (red) traces for three example ROIs. EV: explained ROI variance in test set.

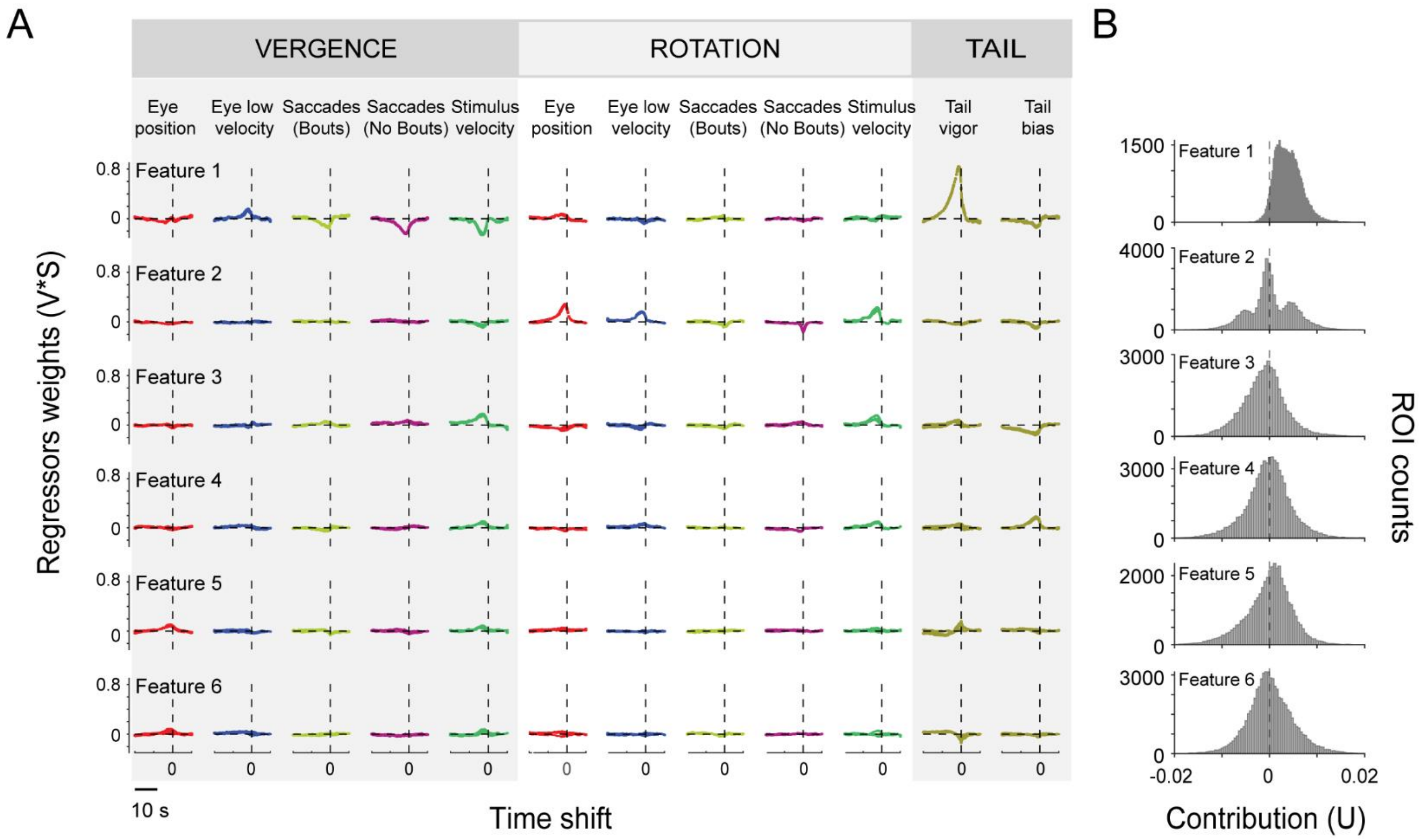


**Figure S6. The best model for the example fish in Figure 4 has six features. (A)** Feature traces scaled by their contribution to population activity (V*S; see Methods). For each feature, variables are color-coded and grouped into vergence, rotation and tail variables, as in Figure 4D. **(B)** Contribution of new regressors associated with features 1 to 6 to the activity of ROIs in the population (N=43622 cubes). Note the highly non-Gaussian distribution of the first two new regressors.

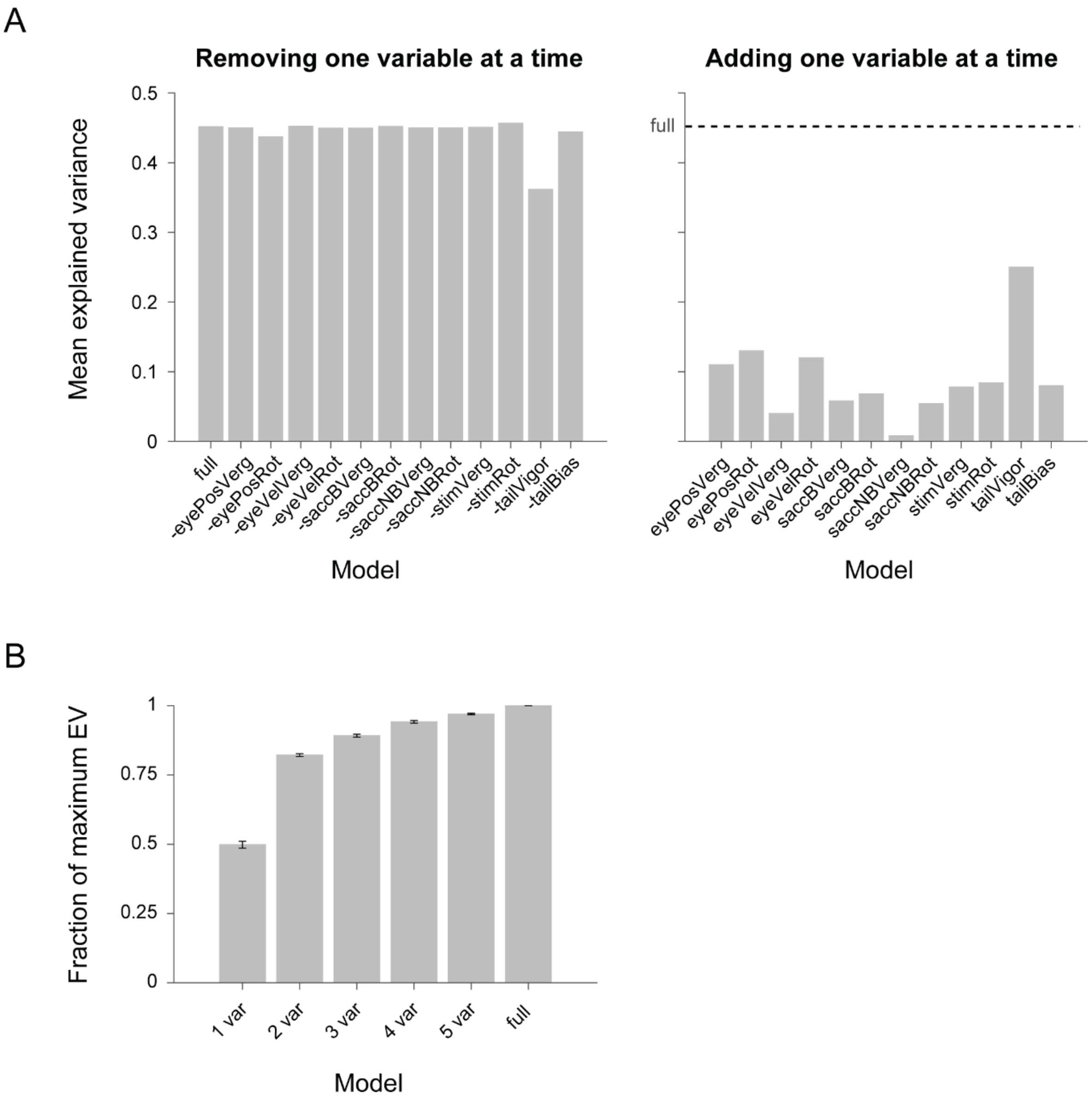


**Figure S7. Variable selection. (A)** Backwards stepwise (left panel) and forward stepwise (right panel) variable selection. Mean explained variance across ROIs of an example fish, when removing one variable at a time (left) or in models with only one variable (right). Note that in both cases, tail vigor has the largest impact on model performance. Eye Position Rotation also affects model performance, although to a lesser degree. **(B)** Fraction of the maximum explained variance (of the full model) for models with one, two, three, four, and five variables. The first and second variables added to the model produce the biggest jump in performance. Adding more variables produces only small increases in the variance explained by the model. Mean ± SEM (N=6 fish). Note that for different fish, the variables incorporated may not be the same in each model.

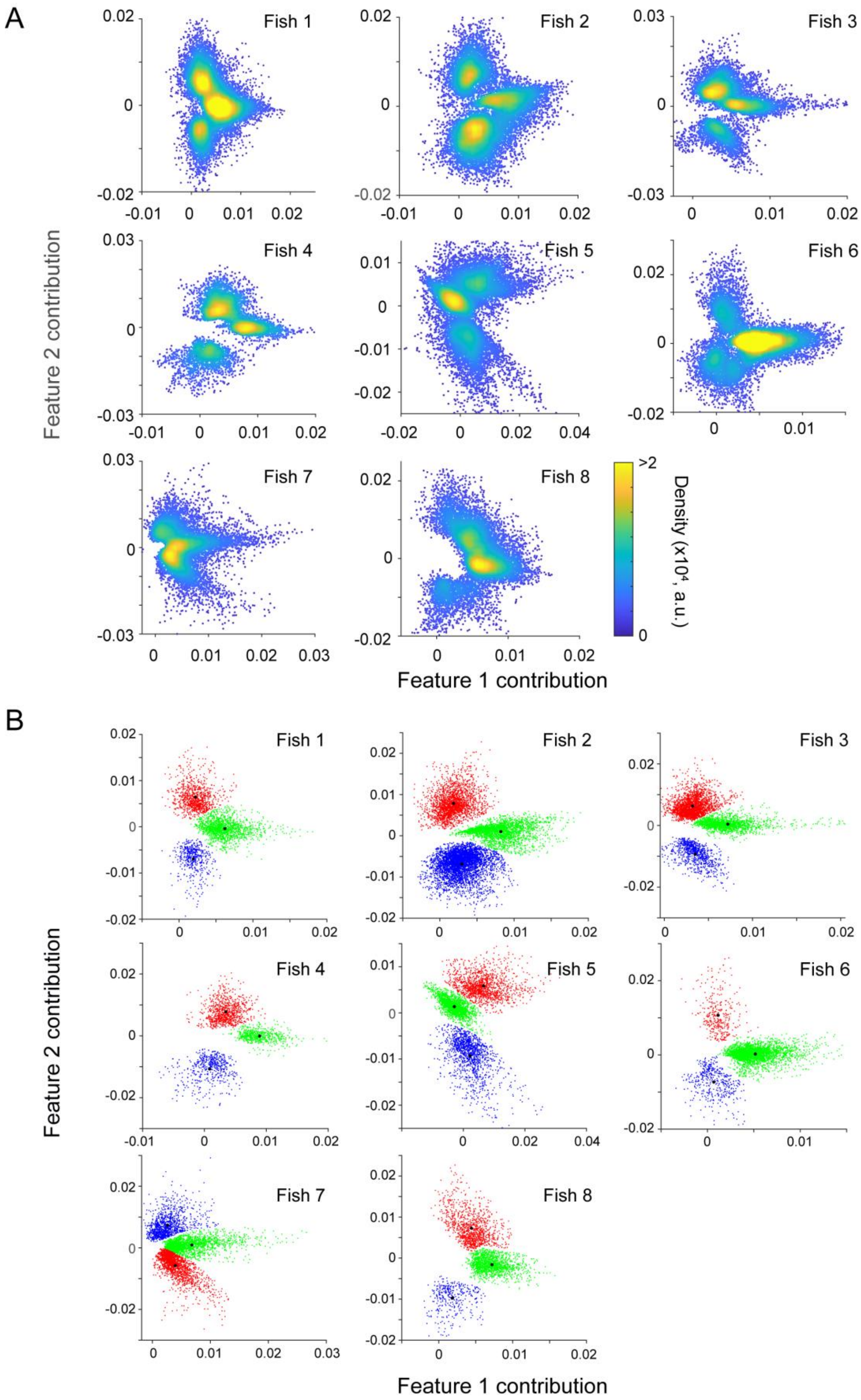
A
Fish 1
Fish 2
Fish 3
Fish 4
Fish 5
Fish 6
Fish 7
Fish 8
Feature 2 contribution
Feature 1 contribution
Density (x10⁴, a.u.)
>2
0
B
Fish 1
Fish 2
Fish 3
Fish 4
Fish 5
Fish 6
Fish 7
Fish 8
Feature 2 contribution
Feature 1 contribution

**Figure S8. Equivalent clusters are found across all imaged fish. (A)** Scatter-density plots for the contributions associated with features 1 and 2 in eight different fish. **(B)** ROIs are assigned to clusters in feature contribution space. ROIs are randomly downsampled to aid visualization in the scatter plot. Black asterisks mark the clusters centroids.

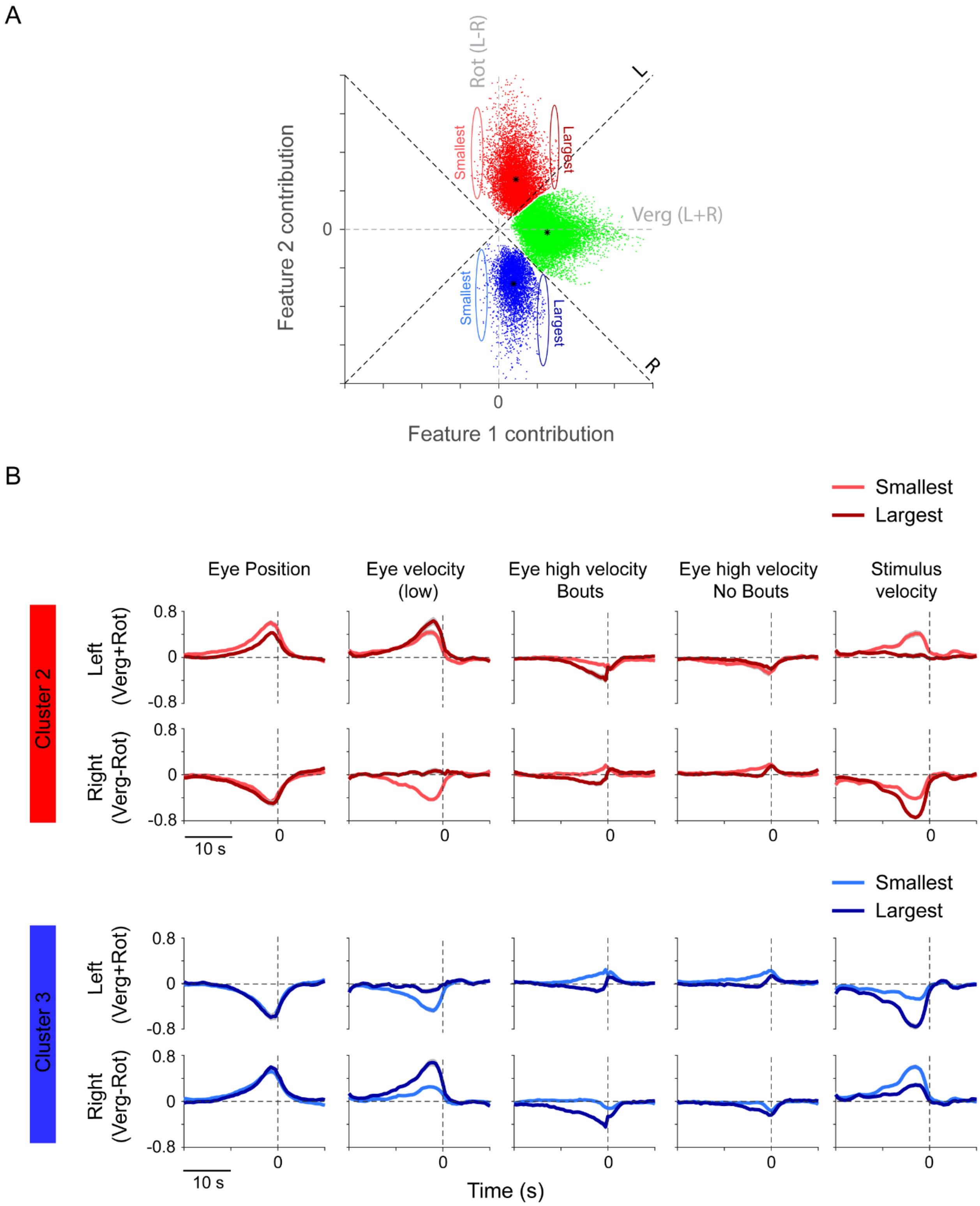
A
Rot (L-R)
L
R
Verg (L+R)
Smallest
Largest
Smallest
Largest
Feature 2 contribution
Feature 1 contribution
0
0
B
Smallest
Largest
Eye Position
Eye velocity (low)
Eye high velocity Bouts
Eye high velocity No Bouts
Stimulus velocity
Cluster 2
Left (Verg+Rot)
Right (Verg-Rot)
0.8
0
-0.8
10 s
Smallest
Largest
Cluster 3
Left (Verg+Rot)
Right (Verg-Rot)
10 s
Time (s)

**Figure S9. Responses of ROIs in the rotation clusters follow a continuum, combining left and right eye information to different extents. (A)** ROIs cluster in feature contribution space. The horizontal axis corresponds to the contribution of the vergence feature (feature 1), while the vertical axis corresponds to the contribution of the rotation feature (feature 2). In the ‘rotation’ clusters (cluster 2, red; cluster 3, blue), ROIs fall in a continuum along the horizontal axis according to the contribution of the vergence feature to their activity. We looked at ROIs at the two extremes, with the smallest and largest contributions of the vergence feature (colored ellipses). **(B)** Representation of each variable along the left and right axis (as a result of summing/subtracting the vergence and rotation kernels, respectively). Each trace is the grand average across 7 fish (±SEM). For each fish, we averaged the kernels of five ROIs with a small contribution of feature 1 (‘smallest’), and five ROIs with a large contribution of feature 1 (‘largest’).

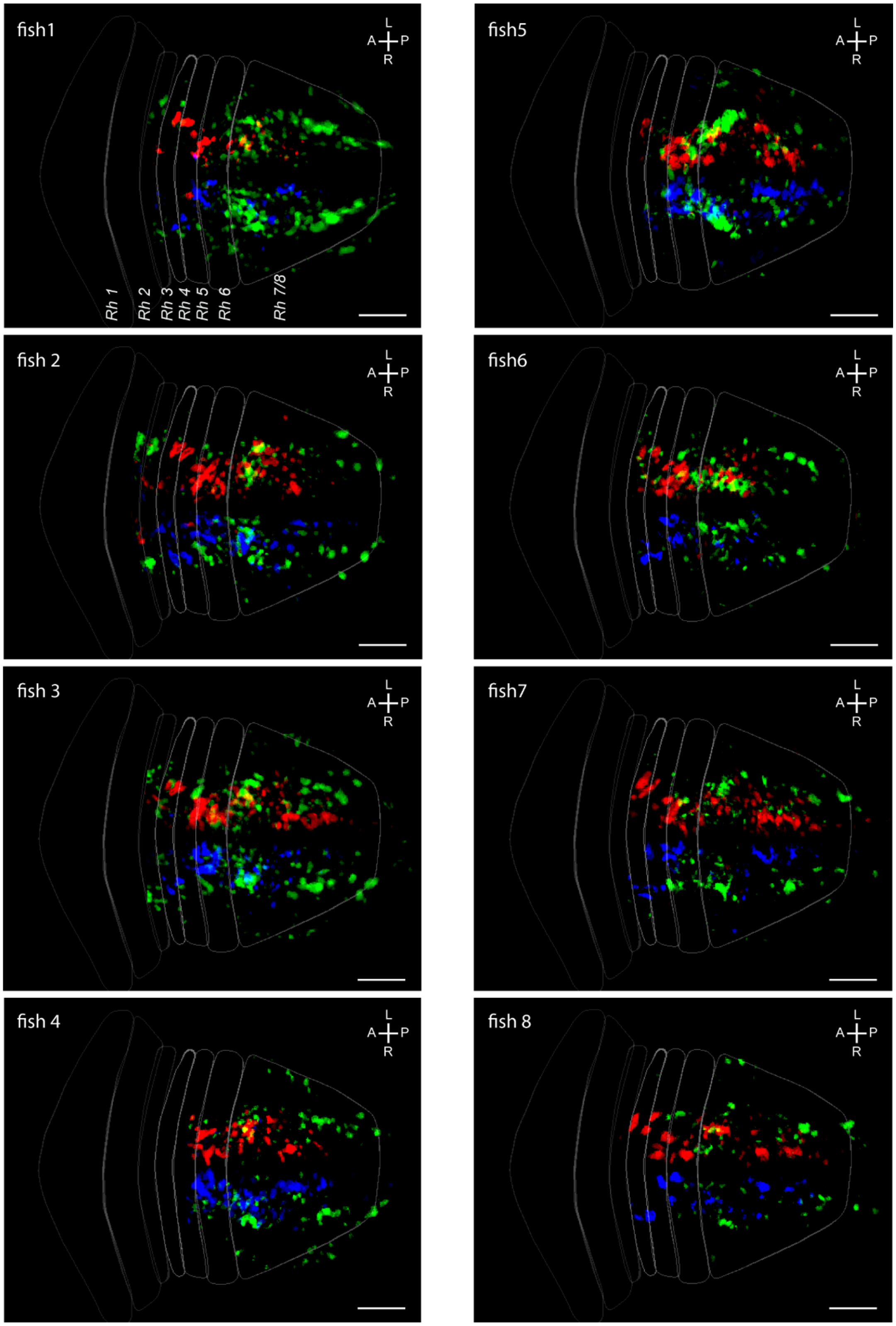
fish1
L
A
P
R
Rh 1
Rh 2
Rh 3
Rh 4
Rh 5
Rh 6
Rh 7/8
fish5
fish 2
fish6
fish 3
fish7
fish 4
fish 8

**Figure S10. ROIs belonging to the three clusters have a stereotyped spatial distribution.** Spatial distribution of ROIs assigned to equivalent clusters in eight different fish (these maps are registered and averaged in Figure 6). Maximum projections of 3D stacks. Rhombomeres contours correspond to Z-brain masks (Randlett et al., 2015). Note that imaged regions are not identical (for instance, for fish 4 the imaging area is a bit more caudal).

### Supplementary movies

**Movie S1.** Visual stimulus. The stimulus was projected on a screen underneath the fish. The velocity of the stimuli on the two hemifield was modulated independently.

**Movie S2.** Anatomical distribution of ROIs belonging to the three clusters. Average map across eight fish.

**Table 1.** List of regressors used for correlation analysis.

| | |
|---|---|
| 1 | left eye position raw temporal positive' |
| 2 | left eye position (temporal positive) threshold:-25' |
| 3 | left eye position (temporal positive) threshold:-20' |
| 4 | left eye position (temporal positive) threshold:-15' |
| 5 | left eye position (temporal positive) threshold:-10' |
| 6 | left eye position (temporal positive) threshold:-5' |
| 7 | left eye position (temporal positive) threshold:0' |
| 8 | left eye position (temporal positive) threshold:5' |

| | |
|---|---|
| 9 | left eye position (temporal positive) threshold:10' |
| 10 | left eye position (temporal positive) threshold:15' |
| 11 | left eye position (temporal positive) threshold:20' |
| 12 | left eye position (temporal positive) threshold:25' |
| 13 | left eye position (nasal positive) threshold:-25' |
| 14 | left eye position (nasal positive) threshold:-20' |
| 15 | left eye position (nasal positive) threshold:-15' |
| 16 | left eye position (nasal positive) threshold:-10' |
| 17 | left eye position (nasal positive) threshold:-5' |
| 18 | left eye position (nasal positive) threshold:0' |
| 19 | left eye position (nasal positive) threshold:5' |
| 20 | left eye position (nasal positive) threshold:10' |
| 21 | left eye position (nasal positive) threshold:15' |
| 22 | left eye position (nasal positive) threshold:20' |
| 23 | left eye position (nasal positive) threshold:25' |
| 24 | right eye position temporal positive' |
| 25 | right eye position (temporal positive) threshold:-25' |
| 26 | right eye position (temporal positive) threshold:-20' |
| 27 | right eye position (temporal positive) threshold:-15' |
| 28 | right eye position (temporal positive) threshold:-10' |
| 29 | right eye position (temporal positive) threshold:-5' |
| 30 | right eye position (temporal positive) threshold:0' |
| 31 | right eye position (temporal positive) threshold:5' |
| 32 | right eye position (temporal positive) threshold:10' |
| 33 | right eye position (temporal positive) threshold:15' |
| 34 | right eye position (temporal positive) threshold:20' |
| 35 | right eye position (temporal positive) threshold:25' |
| 36 | right eye position (nasal positive) threshold:-25' |
| 37 | right eye position (nasal positive) threshold:-20' |
| 38 | right eye position (nasal positive) threshold:-15' |
| 39 | right eye position (nasal positive) threshold:-10' |

| | |
|---|---|
| 40 | right eye position (nasal positive) threshold:-5' |
| 41 | right eye position (nasal positive) threshold:0' |
| 42 | right eye position (nasal positive) threshold:5' |
| 43 | right eye position (nasal positive) threshold:10' |
| 44 | right eye position (nasal positive) threshold:15' |
| 45 | right eye position (nasal positive) threshold:20' |
| 46 | right eye position (nasal positive) threshold:25' |
| 47 | left eye velocity raw temporal positive' |
| 48 | left eye velocity filtered (H+L) temporal positive' |
| 49 | left eye velocity temporal (positive) rectified' |
| 50 | left eye velocity nasal (positive) rectified' |
| 51 | left eye velocity raw LOW temporal positive' |
| 52 | left eye velocity rectified LOW temporal positive' |
| 53 | left eye velocity rectified LOW nasal positive' |
| 54 | left eye velocity raw HIGH temporal positive' |
| 55 | left eye velocity rectified HIGH temporal positive' |
| 56 | left eye velocity rectified HIGH nasal positive' |
| 57 | saccade left eye' |
| 58 | saccade left eye LEFTWARD' |
| 59 | saccade left eye RIGHTWARD' |
| 60 | right eye velocity raw temporal positive' |
| 61 | right eye velocity filtered (H+L) temporal positive' |
| 62 | right eye velocity temporal positive rectified' |
| 63 | right eye velocity nasal positive rectified' |
| 64 | right eye velocity raw LOW temporal positive' |
| 65 | right eye velocity rectified LOW temporal positive' |
| 66 | right eye velocity rectified LOW nasal positive' |
| 67 | right eye velocity raw HIGH temporal positive' |
| 68 | right eye velocity rectified HIGH temporal positive' |
| 69 | right eye velocity rectified HIGH nasal positive' |
| 70 | saccade right eye' |

| | |
|---|---|
| 71 | saccade right eye LEFTWARD' |
| 72 | saccade right eye RIGHTWARD' |
| 73 | saccade either eye, either direction' |
| 74 | left - right eye velocity' |
| 75 | left + right eye velocity' |
| 76 | left - right low eye velocity' |
| 77 | left + right low eye velocity' |
| 78 | left - right high eye velocity' |
| 79 | left + right high eye velocity' |
| 80 | left + right eye temporal position' |
| 81 | left - right eye temporal positions' |
| 82 | left + right eye temporal position rectified' |
| 83 | left + right eye nasal position rectified' |
| 84 | left - right eye temporal position rectified' |
| 85 | left - right eye nasal position rectified' |
| 86 | left eye position + velocity NO saccades rectified temporal' |
| 87 | left eye position + velocity NO saccades rectified nasal' |
| 88 | right eye position + velocity NO saccades rectified temporal' |
| 89 | right eye position + velocity NO saccades rectified nasal' |
| 90 | left eye acceleration (veloc increase) raw temporal positive' |
| 91 | left eye acceleration (all) raw temporal positive' |
| 92 | left eye acceleration nasal (positive) rectified' |
| 93 | left eye acceleration temporal (positive) rectified' |
| 94 | right eye acceleration (veloc increase) raw temporal positive' |
| 95 | right eye acceleration (all) raw temporal positive' |
| 96 | right eye acceleration nasal (positive) rectified' |
| 97 | right eye acceleration temporal (positive) rectified' |
| 98 | Left eye 1/4 position + 3/4 velocity no sacc (temporal positive)' |
| 99 | Left eye 1/2 position + 1/2 velocity no sacc (temporal positive)' |
| 100 | Left eye 3/4 position + 1/4 velocity no sacc (temporal positive)' |
| 101 | Right eye 1/4 position + 3/4 velocity no sacc (temporal positive)' |

| | |
|---|---|
| 102 | Right eye 1/2 position + 1/2 velocity no sacc (temporal positive)' |
| 103 | Right eye 3/4 position + 1/4 velocity no sacc (temporal positive)' |
| 104 | left eye stimulus velocity raw temporal positive' |
| 105 | left eye stimulus velocity rectified nasal' |
| 106 | left eye stimulus velocity rectified temporal positive' |
| 107 | left eye stimulus speed' |
| 108 | left eye retinal slip' |
| 109 | left eye retinal slip rectified nasal' |
| 110 | left eye retinal slip rectified temporal' |
| 111 | left eye relative stimulus speed' |
| 112 | right eye stimulus velocity raw temporal positive' |
| 113 | right eye stimulus velocity rectified nasal (countCLK)' |
| 114 | right eye stimulus velocity rectified temporal (CLK)' |
| 115 | right eye stimulus speed' |
| 116 | right eye retinal slip' |
| 117 | right eye retinal slip rectified nasal (countCLK)' |
| 118 | right eye retinal slip rectified temporal (CLK)' |
| 119 | right eye relative stimulus speed' |
| 120 | left + right stim veloc' |
| 121 | left - right stim veloc' |
| 122 | l. eye clk - r eye clk (rect)' |
| 123 | r. eye clk - l eye clk' |
| 124 | l. eye clk - r eye countclk' |
| 125 | r. eye clk - l eye count' |
| 126 | l. eye countclk - r eye clk' |
| 127 | r. eye countclk - l eye clk' |
| 128 | l. eye countclk - r eye countclk' |
| 129 | r. eye countclk - l eye countclk' |
| 130 | rotation CLK (l. eye nasal & r eye temp)' |
| 131 | stim diverge (l. eye temp & r eye temp)' |
| 132 | stim converge (l. eye nasal & r eye nasal)' |

| | |
|---|---|
| 133 | rotation countCLK (l. eye temp & r eye nasal)' |
| 134 | Relative veloc l. eye CLK - r eye CLK (rect)' |
| 135 | Relative veloc r. eye CLK - l eye CLK' |
| 136 | Relative veloc l. eye CLK - r eye countCLK' |
| 137 | Relative veloc r. eye CLK - l eye countCLK' |
| 138 | Relative vel.l. eye countCLK - r eye CLK' |
| 139 | Relative vel r. eye countCLK - l eye CLK' |
| 140 | Relative vel l. eye countCLK - r eye countCLK' |
| 141 | Relative vel r. eye countCLK - l eye countCLK' |
| 142 | Relative l. eye nasal & r eye temp (right rotation)' |
| 143 | Relative l. eye temp & r eye temp (divergence)' |
| 144 | Relative l. eye temp & r eye nasal (convergence)' |
| 145 | Relative l. eye temp & r eye nasal (left roation)' |
| 146 | Tail position' |
| 147 | Tail bouts (stdev)' |
| 148 | Tail vigor (rolling stdev)' |
| 149 | forward' |
| 150 | Heading' |
| 151 | Heading change' |
| 152 | Tail bias positive leftwards' |
| 153 | left turn' |
| 154 | right turn' |
| 155 | fwd swim' |
